# The single-berry metabolomic clock paradigm reveals new stages and metabolic switches during grapevine berry development

**DOI:** 10.1101/2024.07.06.602344

**Authors:** Flora Tavernier, Stefania Savoi, Laurent Torregrosa, Philippe Hugueney, Raymonde Baltenweck, Vincent Segura, Charles Romieu

## Abstract

- Asynchronous development of berries causes metabolic chimerism in usual samples. We thus revisited the developmental changes in the metabolome of the *Vitis vinifera* single berries from anthesis to over-ripening.
- A dataset of 9,256 ions obtained by non-targeted ultra-performance liquid chromatography coupled to high-resolution mass spectrometry was submitted to an analysis workflow combining classification and dimension reduction tools, to reveal the dynamics of metabolite composition without phenological *a priori*.
- This approach led to a metabolome-based definition of developmental stages, as well as the clustering of metabolites into 12 specific kinetic patterns. The single berry intrinsic metabolomic clock alleviates constitutive asynchronicity biases in the usual combination of phenological scales and observer clock. Such increase in temporal resolution enabled the identification of metabolite clusters annunciative of the onset of ripening since the herbaceous plateau. In particular, these clusters included transient lipidic changes and the start of ABA accumulation. We also highlighted a cluster of stilbenes that accumulate after sugar loading stops, during fruit shriveling.
- This non-targeted approach enables a more precise and unbiased characterization of grapevine berry development through the metabolomic clock paradigm. The discovery of new metabolic milestones of berry development paves the way towards an unbiased assessment of berry physiological stages.

## Introduction

By providing both seed protection and dissemination to Angiosperms (Seymour et al., 2013), the apparition of the fruit stands as a pivotal leap in the evolutionary history of plants and their related animal vectors. Fleshy fruits shift from an immature seed protective organ, in green stage, to a strongly attractive one during ripening, when mature seeds become resistant enough to be disseminated. This key transition includes typical metabolic shifts affecting both primary metabolites serving as major *osmoticum*, from organic acids to energy-rewarding soluble sugars, and secondary metabolites, from astringent and antinutritive tannins to attractive anthocyanins and aromas (Gillaspy et al., 1993). Most fleshy fruits undergo these developmental transformations regardless of their climacteric or non-climacteric status. However, by contrast with climacteric fruit which rapidly ripen when triggered by the ethylene hormone following long periods of post-harvest storage, non-climacteric fruits do not store significant starch reserve during green stage, and are thus forced to ripen on the plant, in real time with phloem unloading of soluble sugars, making them particularly sensitive to environmental conditions during this period (Giovannoni, 2004).

Among non-climacteric fruits, grapevine (*Vitis vinifera*) is one of the world’s most important, cultivated to produce table grapes, wine, juice and other products. Grape phenology was described in two main stages, following a double sigmoid growth curve (Coombe and Hale, 1973; CooMbe and McCarthy, 2000). The first one, also called green stage, starts with intense cell divisions and expansion, resulting from the vacuolar accumulation of malic and tartaric acids as major *osmoticum* (Ojeda et al., 1999; Terrier and Romieu, 2001). Condensed tannins are synthesized at the beginning of this stage (Ollé et al., 2011). These phenolic compounds are important for wine organoleptic qualities (Del-Castillo-Alonso et al., 2021). The second stage, also called ripening, starts with an abrupt softening of the berries indicating the sudden induction of sugar accumulation. Such accumulation involves an accelerated phloem discharge that is accompanied by a significant respiration of malic acid (Coombe, 1992; Shahood et al., 2020). By the time sugars reach their peak, berry volume has doubled due to the parallel import of water. The berries then concentrate all solutes as they shrivel, but no longer import sugars (Castellarin et al., 2007a; Daviet et al., 2023; Savoi et al., 2021). The transition between green and ripening stages is commonly called “veraison” and refers to the start of anthocyanidins accumulation, responsible for the color change in red-skinned grape varieties (Lund and Bohlmann, 2006). By extension, this also indicates the softening and the initiation of sugar loading, slightly before color change, as elucidated on single berries (Bigard et al., 2019; Daviet et al., 2023).

Research on grape development is widely documented, but the division into distinct stages varies between studies (Bigard et al., 2019; Castellarin et al., 2007a; Rogiers et al., 2017; Savoi et al., 2021; Shahood et al., 2020). Stage characterisation relies on the measurement of the main primary metabolites (such as sugars, malic and tartaric acids), berry coloration, softness and growth. Omics technologies have enhanced the exhaustivity of developmental characterization at both the transcriptomic (Cabral et al., 2023; Castellarin and Di Gaspero, 2007; Cramer et al., 2014; Goes da Silva et al., 2005; Savoi et al., 2021; Tornielli et al., 2023) and metabolomic (Bonada et al., 2013; Duan et al., 2019; Leng et al., 2021; Nicolas et al., 2024; Ollé et al., 2011) levels. Furthermore, some studies have adopted an integrative approach that combines both transcriptomic and metabolomic analyses, leading to the identification of genes that could be activated at specific developmental stages and of molecules that could signal or mark stages beyond those defined by growth, primary metabolism and anthocyanins (Fasoli et al., 2018, 2012). However, most of these traits vary continuously during development, making the discretization of development beyond the two main stages somewhat arbitrary.

Research on berry development usually focuses on successives sets of mixed berries in order to smooth their marked heterogeneity at the plot level. Indeed, the berries exhibit both intra and inter cluster developmental asynchrony, ripening both at different rates and dates (Daviet et al., 2023; Shahood et al., 2020). Classical grapevine phenological scales explicitly refers to the median of the population (Coombe, 1995; Lorenz et al., 1995), but it was only recently pointed out that smoothing asynchronous berries obviously generates developmental chimera fundamentally incompatible with the identification of physiological stages as pure metabolic units (Bigard et al., 2019; Shahood et al., 2020). Single berry sampling and characterization have enabled to revisit the flow of water and primary metabolites in depth, to gather quantitative and molecular arguments on the origin of the sugar/acid relationship in grapes, and to shed light on the organization and energetics of the unloading of sugars in the berry. Only studies on single berries have made it possible to establish that softening, initiation of sugar storage, growth resumption and coloration mentioned above occur step by step in this order (Bigard et al., 2019; Savoi et al., 2021; Shahood et al., 2020), while they rather seemed to occur simultaneously on sets of mixed berries (Fasoli et al., 2018).

Recently, new methods have been developed to analyze developmental processes using pipelines combining statistical methods to process omics data. For example, the use of unsupervised learning methods, based on dimension reduction and analysis of distances between samples, has already made it possible to define a new grape phenology scale based on transcriptomics (Tornielli et al., 2023). The present study combines multivariate analyses with untargeted metabolomics on single berries to revisit and refine the phenological stages of grape development. We show that metabolome-wide analyses provide a better understanding of berry development, identifying new metabolites, markers of key steps of berry growth and ripening.

## Material and Methods

### Experimental design

Single berries were sampled by Savoi et al. (2021) in the years 2018 and 2019 on *Vitis vinifera* cv. Syrah within the experimental vineyard Pierre Galet of Institut Agro Montpellier (France). Plants established in 2000, were grafted onto SO4 rootstock and irrigated to avoid severe water deprivation. Throughout the growth cycle, the plants received regular phytosanitary sprayings to limit fungus diseases.

Berries were sampled in accordance with the double sigmoidal growth pattern, as identified through recurring photographic observations and quantification of sugars and organic acids (Savoi et al., 2021). This framework delineated eleven temporal waypoints in the developmental sequence, called « expert stages » for this study and encompassing the green growth phase (G1, G2, and G3), the green lag period (L4 and L5), the onset of ripening (characterized by the softening phase, S6 and S7), the ripening phase (R8, R9, and R10), culminating to the shriveling stage (Sh11) (Savoi et al., 2023). Systematic sampling was undertaken, with the green stage (from G1 to L4) being sampled in 2019, and ripening one (from L5 to Sh11) in 2018. The herbaceous plateau (L4 and L5), allowed to interconnect the two temporal domains.

Berries were frozen without pedicel and seeds in liquid nitrogen, before being crushed using a stainless-steel ball mill (Retsch MM400, Verder Scientific, Inc., Newtown, US). Subsequently, the 153 frozen berry powders underwent a lyophilization for 72h (Cryotec pilot freeze-dryer, Cryotec, Lunel, France), before being analyzed by VIS-NIR reflectance spectroscopy using a LabSpec 2500 with an optical probe (Analytical Spectral Devices, Inc., Boulder, CO, US). These spectra allowed the curated selection of 125 samples, using the Kennard-Stone algorithm (Kennard and Stone, 1969), to ensure parity in berry quantities among stages (Table **1**).

**Table 1.** Repartition of the 125 samples analyzed by UHPLC.

|  | Green |  |  |  |  | Ripening |  |  |  |  | Over-ripening |
| --- | --- | --- | --- | --- | --- | --- | --- | --- | --- | --- | --- |
|  | Growth |  |  | Lag |  | Softening |  | Colouring |  |  | Shrivelling |
| Expert stages | G1 | G2 | G3 | L4 | L5 | S6 | S7 | R8 | R9 | R10 | Sh11 |
| Year of sampling | 2019 | 2019 | 2019 | 2019 | 2018 | 2018 | 2018 | 2018 | 2018 | 2018 | 2018 |
| Harvested samples | 4 | 26 | 28 | 19 | 8 | 9 | 9 | 11 | 14 | 12 | 13 |
| Sample selected | 4 | 14 | 14 | 17 | 8 | 9 | 9 | 11 | 14 | 12 | 13 |

### Metabolomic analyses using ultra high-performance liquid chromatography coupled to high-resolution mass spectrometry (UHPLC-HRMS)

Metabolites were extracted using 20 µL methanol per mg of lyophilized sample, containing 5 µg/mL of chloramphenicol as an internal standard. The extract was sonicated for 10 minutes, before centrifugation at 13000 g at 10°C for 10 minutes. Quality control sample (QC) was prepared by mixing an equal volume of each sample to evaluate the stability of the equipments (retention time and area) and the repeatability of metabolites detection in all of the samples. Supernatants were analyzed as described previously (Rodrigues et al., 2023), with some modifications. Metabolomic analyses were performed using a Vanquish Flex binary UHPLC system (Thermo Scientific, Waltham, MA) equipped with a diode array detector (DAD). Chromatographic separation was performed on a Nucleodur C18 HTec column (150 × 2 mm, 1.8 μm particle size; Macherey-Nagel, Duren, Germany) maintained at 30°C. The mobile phase consisted of acetonitrile/formic acid (0.1%, v/v) (eluant A), and water/formic acid (0.1%, v/v) (eluant B), at a flow rate of 0.25 mL/min. The gradient elution program was as follows: 0 to 4 min, 80% to 70% B; 4 to 5 min, 70% to 50% B; 5 to 6.5 min, 50% B isocratic; 6.5 to 8.5 min, 0% B; and 8.5 to 10 min, 0% B isocratic. The injected volume of sample was 1 μL. The UHPLC was coupled to an Exploris 120 Q-Orbitrap MS system (Thermo Scientific, Waltham, MA) operated with a heated electrospray ionization source in positive and negative ion modes. The key parameters were as follows: spray voltage, + 3.5 and − 3.5 kV; sheath-gas flow rate, 40 arbitrary units (arb. unit); auxiliary-gas flow rate, 10 arb. unit; sweep-gas flow rate, 1 arb. unit; capillary temperature, 320°C; and auxiliary-gas-heater temperature, 300°C. The scan modes were full MS with a resolution of 60 000 fwhm (at m/z 200) and ddMS2 with a resolution of 60 000 fwhm; the normalized collision energy was 30 V; and the scan range was m/z 85−1200. Internal mass calibration was operated using EASY-IC internal calibration source allowing single mass calibration for full mass range. Data acquisition and processing were carried out with Xcalibur 4.5 and Free Style 1.7 (Thermo Scientific, Waltham, MA), respectively. Analyses were performed in both positive and negative ionization modes, thereby constituting dual data sets for each individual sample.

UHPLC-HRMS raw data were processed using the Compound Discoverer 3.3 software (Thermo Fisher Scientific, Waltham, MA, USA). Data processing for positive and negative modes were performed separately. QC samples were used to verify the repeatability of retention times and signal intensity throughout the data set. Untargeted metabolomics workflows were used for peaks detection, peaks groupement and alignment, and fill in missing peak data. Background compounds found in the blank samples, as well as known environmental contaminants related to plant protection products were filtered out from the data set. Peaks alignment parameters mainly included mass tolerance and retention time (RT), which were set at 5 ppm and at 0.1 min, respectively. Peaks detection was performed using a signal-to-noise ratio (S/N) of 2 and peak intensity thresholds at 10,000. Poorly repeatable ions were filtered out by keeping those with a peak rating greater or equal to 4 in at least 4 samples and by keeping ions with a coefficient of variation (CV) ≤ 30% in all QC samples. Using these settings, 5123 and 4133 ions were obtained in positive and negative modes, respectively. Metabolomics data have been deposited to the EMBL-EBI MetaboLights database (DOI: 10.1093/nar/gkad1045, PMID:37971328) with the identifier MTBLS10572.

### Data analysis and classifications

The high dimensional dataset documenting the evolution of the berry metabolome across the 11 expert documented phenological stages (MS data) was further processed to classify metabolites by similarities in their evolution, using a workflow based on several multivariate analyses (Fig. **1**).

**Figure 1.**
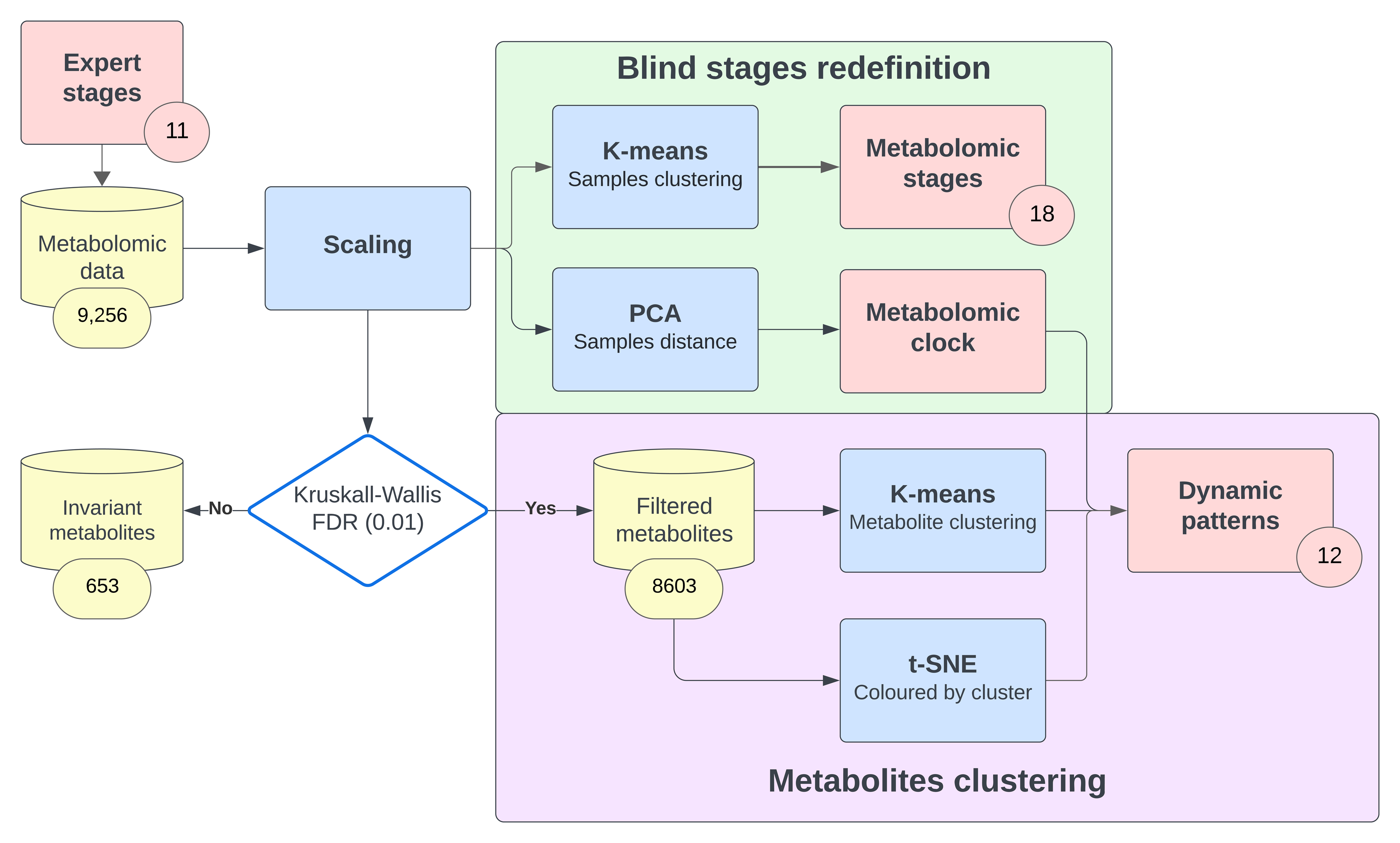
Metabolomic data analysis workflow. The 11 expert stages were defined before our study by Savoi, et al. (2023). The workflow starts with raw metabolomic data, each of the 9,256 detected ions surfaces is initially scaled by its sum over berry development. The normalized data underwent a classification of the samples by the K-means algorithm, in order to identify new metabolomic stages independently from expert stages and observer time. In parallel, the normalized data were submitted to a principal component analysis (PCA). Single berry molecular clock was calculated from the curvilinear distance of samples projected on successive linear regressions, selecting the best principal component (PC) plane for each one. These newly identified stages and clock were then compared to the expert stages. 653 invariant ions were then filtered out using a Kruskal-Wallis test with a false discovery rate of 0.01. The 8,603 remaining ions were then grouped together using K-means on the metabolites, followed by a PCA for each metabolite cluster, enabling the representative dynamics (or “eigen-metabolite”) to be shown. Finally, a t-distributed stochastic neighbor embedding (t-SNE) representation allowed more direct insights on the data structure.

Each peak surface was normalized by their sum of areas measured throughout berry development. Berry samples were classified using the K-means algorithm on the scaled metabolomic matrix to redefine developmental stages, based on variations in metabolite composition. The statistically relevant cluster number was determined using the average silhouette method (Govender and Sivakumar, 2020; Rousseeuw, 1987). The scaled metabolomic matrix was also submitted to a principal component analysis (PCA). The successive samples, projected onto the first three principal components (PC), were spread on a series of linear trajectories separated by abrupt directional changes. Linear regressions were calculated on all possible planes, before selecting the best ones based on their coefficient of determination. The curvilinear distance was then calculated, following an orthogonal projection of each sample on the respective lines, and tracing the progression of samples in the interconnected regressions. Such distance enabled the definition of a continuous metabolomic clock.

Developmentally invariant metabolites were filtered out using the Kruskall-Wallis test with a false discovery rate of 0.01. Subsequently, the metabolites were clustered according to their similarities in developmental profiles, using the K-means algorithm and the average silhouette method. A PCA on the metabolites within each cluster yielded the specific profile of its representative “eigen-metabolite”. Their developmental patterns were finally plotted with respect to the newly defined metabolomic clock. Finally, the filtered MS data were processed with the t-distributed stochastic neighbor embedding (t-SNE) method, yielding a three-dimensional projection of the metabolites. t-SNE was performed using a perplexity parameter of 30 over 1,000 iterations. This approach was designed to preserve the local structures of the data, providing a more accurate and faithful representation than that obtained by PCA, and to validate the use of the K-means method against the actual distribution of the data (Hebra et al., 2021; Olivon et al., 2018; Sorkun et al., 2022).

### Identification of key metabolites

Metabolites presenting the highest correlation with the eigen-metabolite (correlation > 0.8) were selected for identification in each cluster showing interesting dynamics. Ion and metabolite annotations were based on expert analysis of molecular formulae, mass spectra and MS/MS fragmentation patterns in comparison with authentic standards when available and with data from MassBank (https://massbank.eu/MassBank/)), mzCloud (https://www.mzcloud.org/) and Chem-Spider (http://www.chemspider.com/) using the Compound Discoverer 3.3 software. Standards for metabolite identification were purchased from Sigma-Aldrich (Saint-Quentin Fallavier, France) and Extrasynthese (Lyon, France). Identification confidence levels (ICL) established by Schymanski et al. (2014) were applied to score identifications.

## Results

### De novo deciphering of grape developmental stages based on single berry metabolomics

A total of 9,256 ions were detected in the 125 berry samples. The 11 expert stages relied on observations of growth stages, accumulation of primary metabolites (sugars, tartaric and malic acids and potassium) and the softening date. In contrast, the K-means blind analysis of metabolomic data allowed to distinguish 18 developmental stages (Table **2**), henceforth coined “metabolomic stages”. The intrinsic chemical profile of the fruits enabled to break down expert stages into sub categories, offering a more accurate view of berry developmental cycle from early to late stages. For example, the expert stage L4, which brings together transitional samples between the first and second part of the green phase (Fig. **2a**), was subdivided into 5 distinct metabolomic stages (Table **2**), while the L5, G1 and R9 expert stages precisely matched single metabolomic stages (Fig. **2a**). Reciprocally, metabolic stages could also include samples from different expert stages, frequently contiguous, but not only (Table **2**, Fig. **S1**).

**Figure 2.**
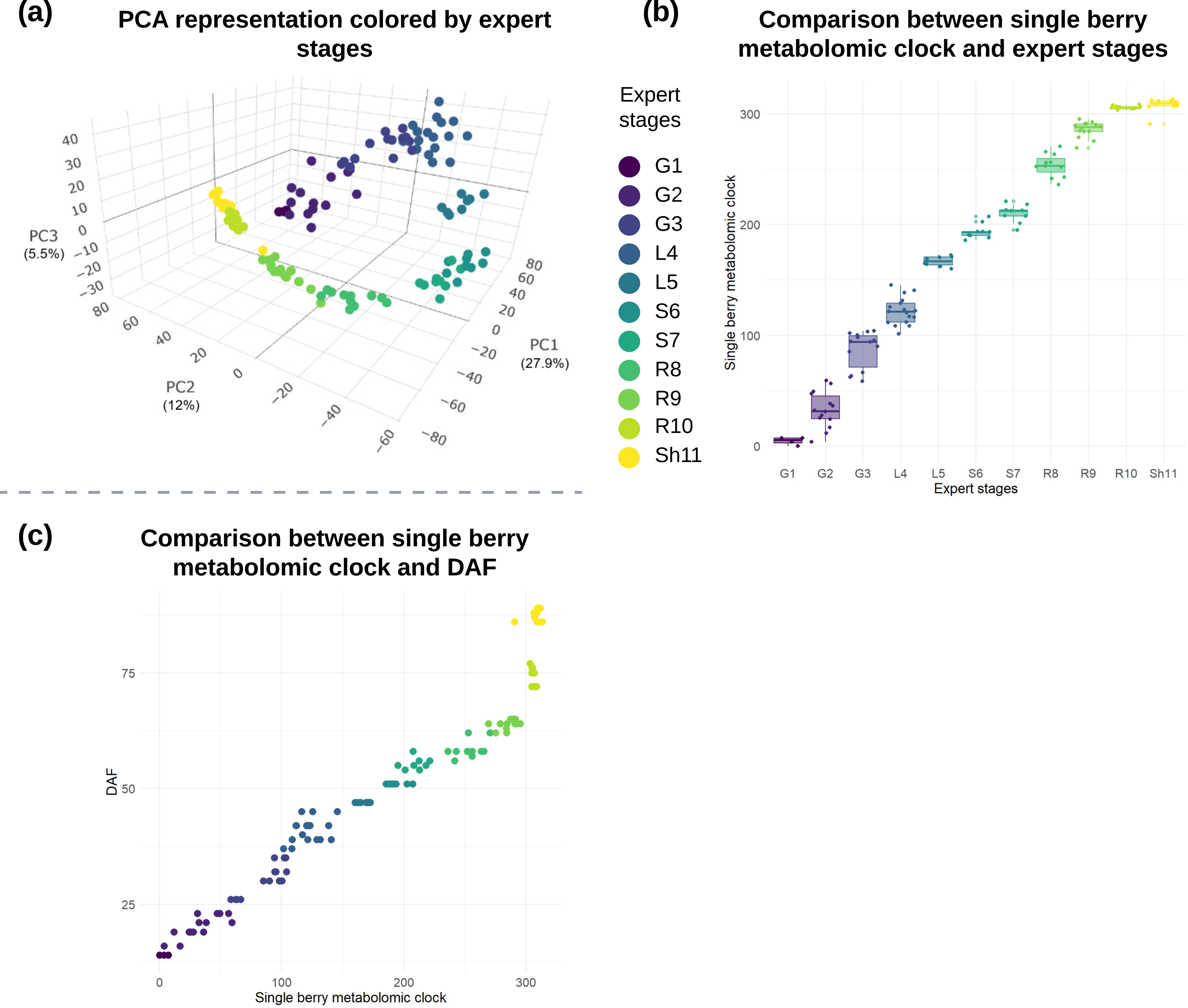
Comparative analysis of berry developmental stages: expert classification single berry metabolic clock and days after flowering (DAF), color-coded by expert stages. (**a**) 3D principal component analysis (PCA) plot displaying berries colored according to expert-defined phenological stages (G = green, L = lag, S = softening, R = ripening, Sh = shriveling). Each color represents a different stage, illustrating the separation and distribution of berries within the multi-dimensional space defined by the first three principal components (PC) of the dataset. (**b**) Scatter plot showing a side-by-side comparison of the metabolomic clock (y-axis) against the expert-defined stages (x-axis). Data points are color-coded by the expert stages. The metabolomic clock was determined by calculating the distances between samples across successives linear regressions of PCA-projected samples trajectories in **a**, providing a continuous scale that contrasts with the discrete categorization of the expert stages. (**c**) Comparison of the metabolomic clock (x-axis) and days after flowering (DAF) (y-axis) and colored by expert stages.

**Table 2.** Relationship between expert stages and K-means determined metabolomic stages. Each cluster represents a stage where berries exhibit similar metabolomic characteristics, G = green phase, L = lag phase, S = softening, R = ripening and Sh = shriveling.

| Expert stages | G1 | G2 | G3 | L4 | L5 | S6 | S7 | R8 | R9 | R10 | Sh11 |
| --- | --- | --- | --- | --- | --- | --- | --- | --- | --- | --- | --- |
| K-means stages | 1 | 1,2,3,4 | 4,5 | 5,6,7,8,9 | 10 | 11,12,13 | 12,13 | 14,15 | 15 | 16,17,18 | 15,17,18 |

In parallel, a “metabolomic clock” has been established by applying linear regressions on PCA-projected sample trajectories, calculating the distances between samples. The 3D PCA projection, where PC 1 to 3 accounted for respectively 27.9%, 12% and 5.5% of the dataset variance, allowed the assessment of the distribution of berries and its comparison with expert stages annotations (Fig. **2a**). The dispersion around the 3D curve decreased from green stage to softening, and became almost non-existent thereafter. According to this projection, the grape development followed 5 main successive trajectories delimited by abrupt changes in direction during L4, S6, R8 and end R9. The “early green stage” (spanning G1 to G3 and some L4 samples) and the “herbaceous plateau stage”, also known as the latent stage (encompassing certain L4 berries and all L5 ones), are distinctly demarcated on the PC2/3 plane. Notably, PC3 clearly distinguished the L4, L5 and S6 expert stages subsequent to sudden changes in berry composition. This segment of the PCA plot is sparsely populated, indicating a potential lack of samples from this stealthy stage. The ripening period itself is subdivided into three trajectories: an “early ripening” group composed of soft and green berries (S6 S7), however these samples harvested a few days apart largely overlap. The subsequent trajectory included R8, R9, and one berry from the Sh11 stage, with a large evolution on PC2. The final trajectory, encompassing R10 and the remaining Sh11 berries, represented the post phloem arrest stage (Savoi et al., 2021) as shown in Fig. **2a**. Fig. **2b** shows that expert stages are less resolutive than metabolic clock, particularly for stages G2, G3 and L4. In contrast, expert stages G1, L5, S6, S7, R10 and Sh11 group berries in a narrower range of metabolomic time. When comparing days after flowering (DAF) and the metabolomic clock (in Fig. **2c**), a fast evolution can be observed inside the L4 expert stage and at the end of maturation. The 50 % flowering date is measured at the scale of the whole cluster and not for each single berry, so it does not take asynchrony into account. It is also important to note that the two sampling seasons (G1 to L4 for 2019 and L5 to Sh11 for 2018, Table **1**) do not appear to be sharply separated on the PCA and graphical representations of Fig. **2**. Altogether, these results clearly show that the metabolomic synchronization of berries outperforms the most severe sorting procedures based on observer sampling time, relative growth and primary metabolites.

### Clustering of the untargeted metabolites using K-means

Clustering single berry metabolites according to their relative amounts in the extracts of 125 single berries revealed 12 major developmental patterns, labeled A to L. These profiles showed 3 major trends: clusters of metabolites that decreased along development (Fig. **3**, profiles A to D), those showing a peak at certain stages (Fig. **4**, profiles E to I) and finally, clusters with a metabolite content that increased along berry development (Fig. **5**, profiles J to L).

**Figure 3.**
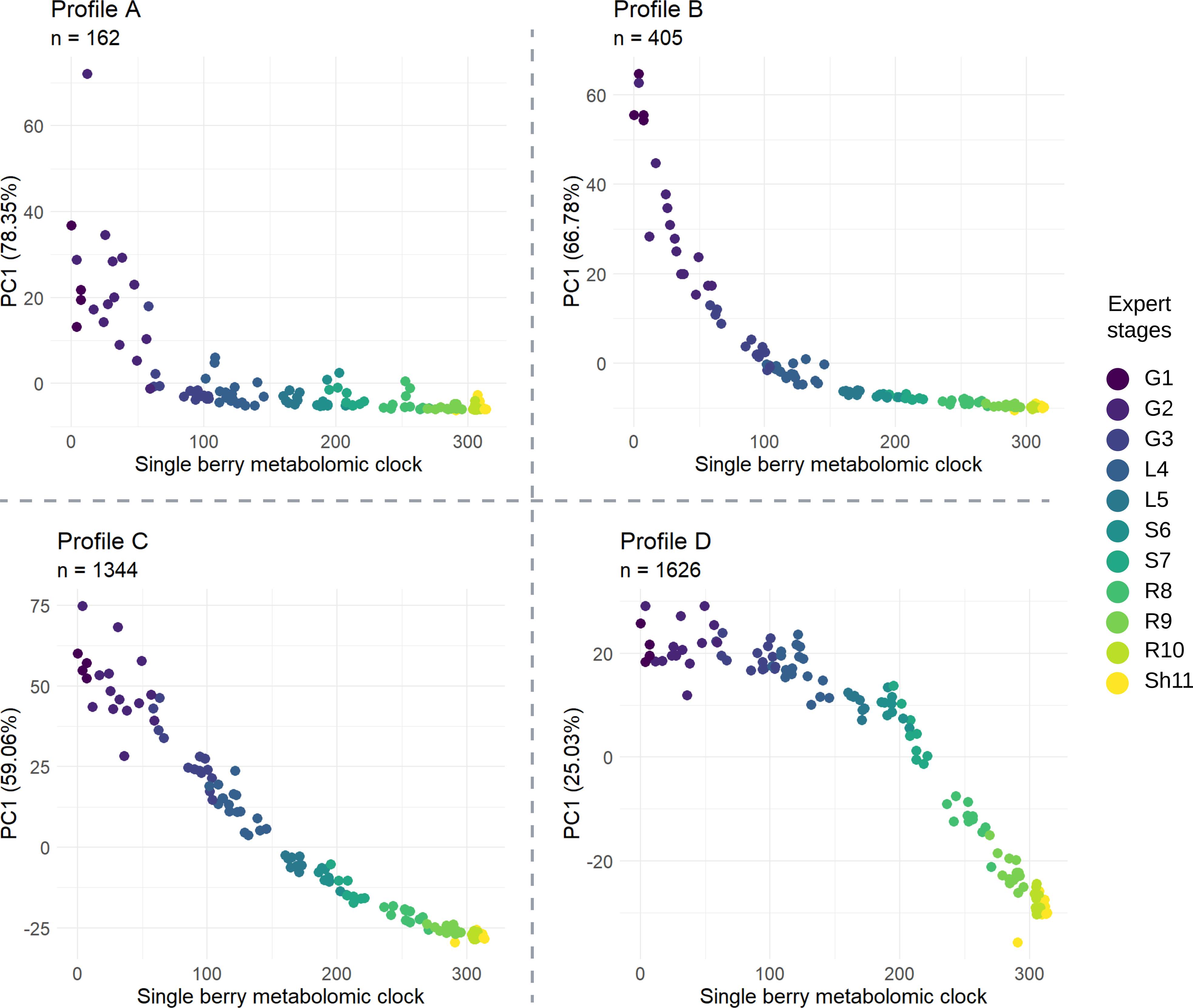
Representative profiles of metabolite clusters decreasing throughout grapevine berry development. n indicates the number of metabolites within each cluster. Each point is a single berry. The y axis represents the first principal component (PC1) from the cluster principal component analysis (PCA) and the x axis is the metabolomic clock. For expert stages description see Table **1**.

**Figure 4.**
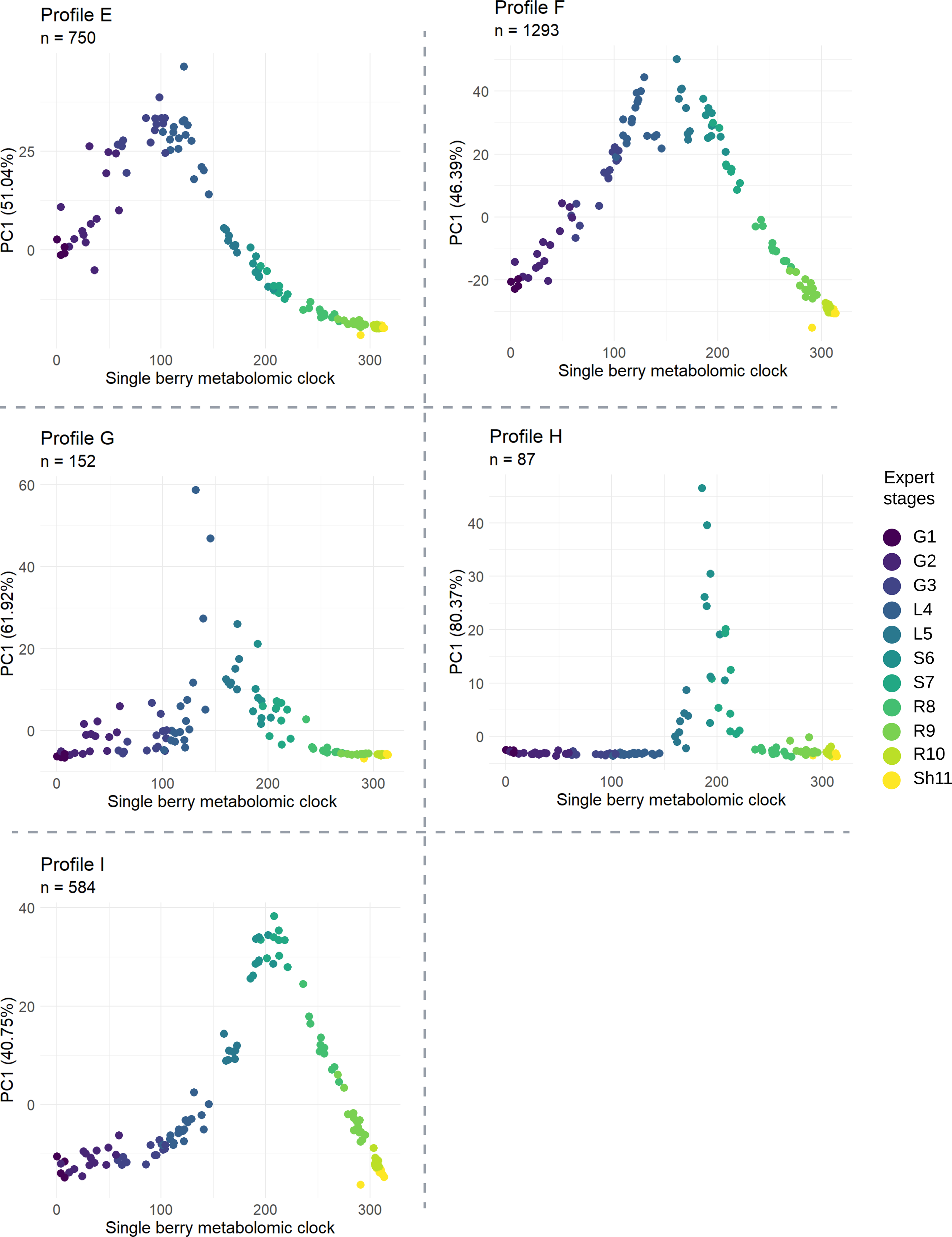
Representative profiles of metabolites that accumulate as a peak at specific stages of the berry development. n indicates the number of metabolites within each cluster. Each point is a single berry. The y axis represents the first principal component (PC1) from the cluster principal component analysis (PCA) and the x axis is the metabolomic clock. For expert stages description see Table **1**.

**Figure 5.**
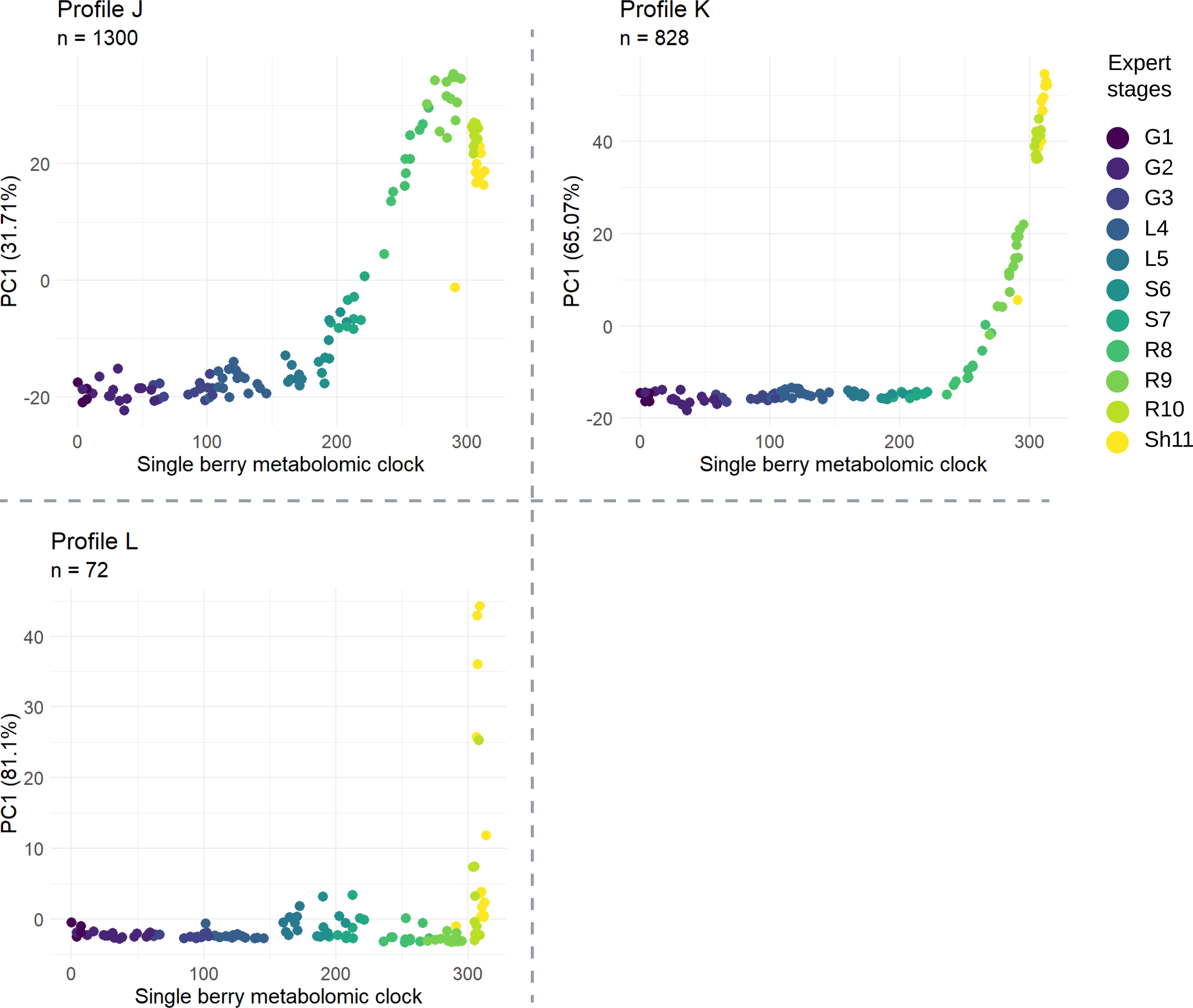
Clusters representative profiles of metabolites showing an accumulation trend during the ripening phase. n value indicates the number of metabolites within each cluster. Each point is a single berry. The y axis represents the first principal component (PC1) from the cluster principal component analysis (PCA) and the x axis is the metabolomic clock. For expert stages description see Table **1**.

The decreasing profiles A, B and C (Fig. **3**) comprised 161, 405 and 1339 metabolites respectively, that could be efficiently summarized with a single PC explaining more than 50% of the intra-cluster variance. In contrast, profile D is distinguished by a larger number of metabolites (1,630), but its PC1 captured a smaller proportion of the total variance (25%). It suggests that this cluster may include more heterogeneous developmental profiles, as underlined by the distribution of correlations between individual metabolite profiles and that of the corresponding eigen-metabolite (Fig. **S2**). The decline of profile A was particularly fast and occurred in the very early green stage, during the G1 and G2 expert stages. Metabolites in profile B decreased continuously through the green stage, from G1 to L4. Profile C exhibited a more gradual decrease over time, starting from a peak at the beginning of the early green stage and declining until the shriveling stage. In contrast, profile D displayed an unusual pattern with a slight decrease during the entire green phase (from the early green stage to the herbaceous plateau) followed by a sharper decline starting at the softening stage and continuing to the shriveling stage.

Figure 4 shows the profiles exhibiting accumulation peaks at different times during berry development. Among these, G and H profiles displayed particularly transient peaks. These profiles contained 146 and 87 metabolites respectively, and could be efficiently summarized with a single PC that explained more than 60% of their variation. Interestingly, the peak in the G profile coincided with the herbaceous plateau phase (L4 and L5), commonly described as a state of metabolic homeostasis. In contrast, profiles E, F and I showed a more gradual accumulation dynamic extending over several stages. These profiles included a higher number of metabolites, 747, 1,296 and 596 respectively, and a lower percentage of variance explained by their first PC by comparison with profiles G and H (Fig. 4, Fig. **S2**). Profile E showed significant accumulation during the early green stage, peaking at the G3 stage and then gradually decreasing. Interestingly, this peak occurred exactly when profile A reached its minimum. Profile F showed a peak during the herbaceous plateau stage (expert stages L4 and L5), marking a transition between the early green stage and the beginning of the ripening. The G profile exhibited a much more abrupt peak compared to profiles E and F, with particular points reaching their maximum at the beginning of the herbaceous plateau stage and then decreasing at the beginning of the softening stage, specifically between stages L4 and S6. The H profile was characterized by an even more pronounced peak, starting at the end of the herbaceous plateau stage (L5) and reaching its maximum during the softening stage (S6 and S7), marking the beginning of grape ripening. At the same time, the I profile also reached its maximum, but with a more gradual increase and decrease, starting at the end of the early green stage (between G3 and L4) and ending at the completion of berry development (Sh11).

Clusters profiles with an ascending shape starting at the S6 expert stage encompassed metabolites specifically accumulated during the berry ripening phase (Fig. 5). Profile J began to accumulate from the start of the softening stage (S6), reaching a peak at the end of the ripening stage, before decreasing during the shriveling stage. Profile K started to increase at the end of the softening stage, from expert stage R8, and reached its maximum at the end of shriveling (expert stage Sh11). Profile J stands out for its large number of metabolites, with 1,297 compounds, and a PC1 that explains only 31% of their variance. On the other hand, profiles K and L contained a smaller number of metabolites, 827 and 72 respectively, and a larger part of their variability was explained by their first PC. The L profile in particular included the largest proportion of metabolites with a correlation of at least 0.8 with the eigen-metabolite (Fig. **S2**). This profile showed a late pattern of accumulation, after phloem arrest and during shriveling (expert stages R10 and Sh11).

Finally, the t-SNE method provides an alternative to PCA for metabolites visualization (Fig. 6). The metabolites separated into two main branches on the 3D graph, reflecting their evolution over time. Early green stage specific patterns (A, B, C and D in shades of blue) are at the upper end of the t-SNE, while dynamics varying during ripening and shriveling stages (J, K and L in shades of red) are at the lower left end of the graph. The dynamics showing accumulation peaks between the early green stage and the ripening stage are in the central part. Transient dynamics observed in clusters A, E, F, G, H, I, J, and L indicate abrupt variations at specific points in berry development, facilitating accurate estimation of developmental stages. In addition, some of these clusters, such as A, H and I, are particularly distinct and clearly separated from the others.

**Figure 6.**
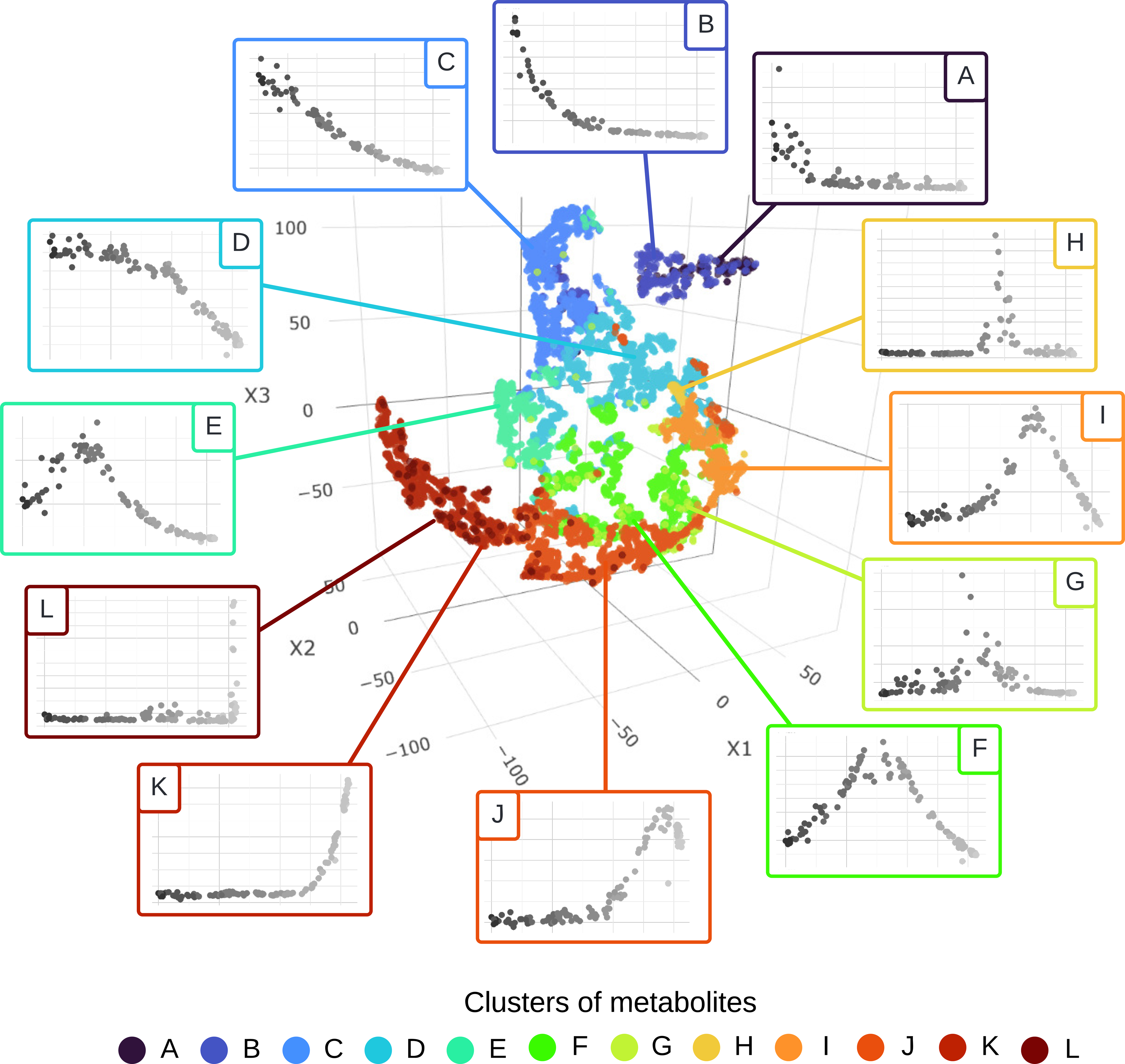
3D t-distributed stochastic neighbor embedding (t-SNE) representation of the untargeted metabolites. Each point represents a metabolite, colored by its K-means cluster with its representative pattern over the metabolomic clock (see details in Fig. **2**, **3** and **4**). Patterns A through D include metabolites consumed in the early green stage. Between E and I, there are noticeable peaks in metabolite accumulation, signaling the stepwise transition from green phase to ripening. Finally, patterns J to L depict the accumulation of metabolites toward the end of the berries’ development. In the 3D t-SNE visualization, these three cluster groups appear to be organized into three distinct branches.

### Metabolite annotation in selected clusters

Among the 12 metabolite clusters, those with the most transient dynamics were chosen for annotation of their metabolites. Profiles A, C (Fig. 3), E, F, H, I (Fig. 4), J, K and L (Fig. 5) were therefore selected, focusing on particularly well classified ions, *i.e.* having a correlation greater or equal to 0.8 with their respective eigen-metabolites. The total number of selected ions was 2,586, and ranged from 60 to 606 for L and C clusters, respectively. Expert analysis of MS and MS/MS data in the 10 selected clusters resulted in the annotation of 483 of the 2,586 ions (Table **S1**), which could be attributed to a total of 117 metabolites, listed in Table **S2**, together with their molecular families. Thus, 19 % of all selected ions could be annotated, this proportion ranging from 7 % (cluster E) to 83 % (cluster L) (Fig. **S3**, Table **S1**).

Cluster A was mainly composed of monomers and small polymers of gallate ester tannins (Fig. **S4**), including confirmed compounds by molecular standard (ICL 1 on the Schymanski et al. (2014) scale) such as epicatechin gallate (ECG) and epigallocatechin gallate (EGCG). Although condensed tannins were predominant in cluster C, particularly procyanidins of types A, B, and C, a few type A procyanidins were also detected in cluster A. Cluster C also contained various flavonoids, including glycosylated derivatives of chalcone and kaempferol, as well as their dihydro-derivatives (ICL 4). Additionally, quercetin glucuronide (ICL 2) was identified in this cluster, along with phenolic acids coupled with tartaric acid, including caftaric, coutaric, and fertaric acids (ICL 1). Malic and citric acids were detected in cluster F (ICL 1). Cluster H encompassed C_18_ and C_20_ lipids, notably linolenic acid (C_18_), confirmed by molecular standard (ICL 1), and its potential derivatives, noted as ICL 2 and 3. Aspartic acid was identified in cluster I (ICL 1), along with polyol sugars and hexose (ICL 2). Cluster J included an anthocyanin, cyanidin glucoside (ICL 1), flavonoids such as derivatives of isorhamnetin and syringetin (ICL 3), the glycosylated form of abscisic acid (ABA-glucoside), as well as glycosylated and esterified coumaric acid and sugar polymers (ICL 1 and 2). Cluster K contained several flavonoids, including derivatives of isorhamnetin and kaempferol (ICL 1 for glycosylated forms and ICL 3 for other derivatives), as well as anthocyanins like myricetin (ICL 2) and malvidin (ICL 1), and contained ABA-glucoside (ICL 2) and proline (ICL 1). Given that ABA-glucosides were present in both clusters J and K, The profile of free ABA was searched for among metabolites with a correlation lower than 0.8 with the eigen-metabolite. Indeed, the free form of ABA had a correlation of 0.75 with the eigen-metabolite of cluster I (Table **S1**). Finally, based on the annotation of the vast majority of its ions (83%), the metabolites in group L belonged almost exclusively to the stilbene family, including isomeric forms and derivatives of resveratrol and piceid (ICL 1), as well as viniferin glucoside (ICL 1).

## Discussion

Present work provides a new perspective on grapevine berry development based solely on its solute composition to establish an internal metabolomic clock. By employing a novel strategy that combines single berry sampling, untargeted metabolomics, multivariate analyses and expert annotation, we provide a much detailed description of the phenological sequence that undergoes grapevine fruit. This approach revealed potentially important groups of metabolites in fruit development. Furthermore, the integration of clustering coupled with t-SNE, applied independently from observer time, led to a more precise and incisive understanding of the grape ripening process.

### Single berry metabolomics reveals an internal clock regulating grape development

The blind statistical analysis of metabolites at single berry level highlighted transient metabolic shifts, which would have been smoothed by averaging asynchronous berries as usually carried out (Bigard et al., 2019; Shahood et al., 2020; Zamboni et al., 2010). In addition, the main solutes that exhibit monotonous changes during green stage or ripening, such as malic acid and sugars, do not allow these phases to be divided into transient stages. We have shown that relying only on variations in fruit intrinsic composition led to a more accurate synchronization of berries than sorting by growth or primary metabolites. The use of PCA to get insights on sample distribution during fruit development has already revealed a common “U” shape pattern throughout the literature, but uncontrolled variations in the real age pyramid of berries inside each sample scatters the data and prevents to detect discontinuities (Dai et al., 2013; Tornielli et al., 2023). Without prior filtering of the data except invariant metabolites, sub periods in the green and ripening stages were clearly resolved on the first 3 PCs, being discriminated by very sudden changes in sample direction vectors, which are necessarily smoothed in average samples. The t-SNE method confirmed the robustness of using metabolites to characterize grape development, organizing them within a developmental continuum. This technique also highlighted isolated groups (clusters A, H, and I) which, although distinct, remained integrated within the developmental flow.

### Profiles are consistent across harvest years

The kinetics of the two sampling years were linked by the lag phase, during which berry growth pauses and which is widely accepted as quite stable during grape development (Coombe, 1976; Matthews et al., 1987; Thomas et al., 2008). Examination of the PCA and representative dynamics revealed a remarkable consistency: the change of vintage did not seem to affect the evolution of the samples. Indeed, the L5 group was not impacted by the sampling year and was equidistant from the other stages (L4 and S6). This continuity was also obvious on representative dynamics, particularly in profiles B, C, D, I, K and L (Fig. **S4**). These results validated our sampling procedure and underlined a certain robustness over two vintages.

### Metabolic clusters reveal transient shifts in berry physiology

Blind, dynamic-based classification of grape metabolites resulted in 12 distinct developmental clusters. Among these, 10 showed relatively transient dynamics, enabling us to trace the developmental history of grapevine berry and propose a metabolic map, which highlights some metabolites and pathways as milestones of grape development (Fig. 7).

**Figure 7.**
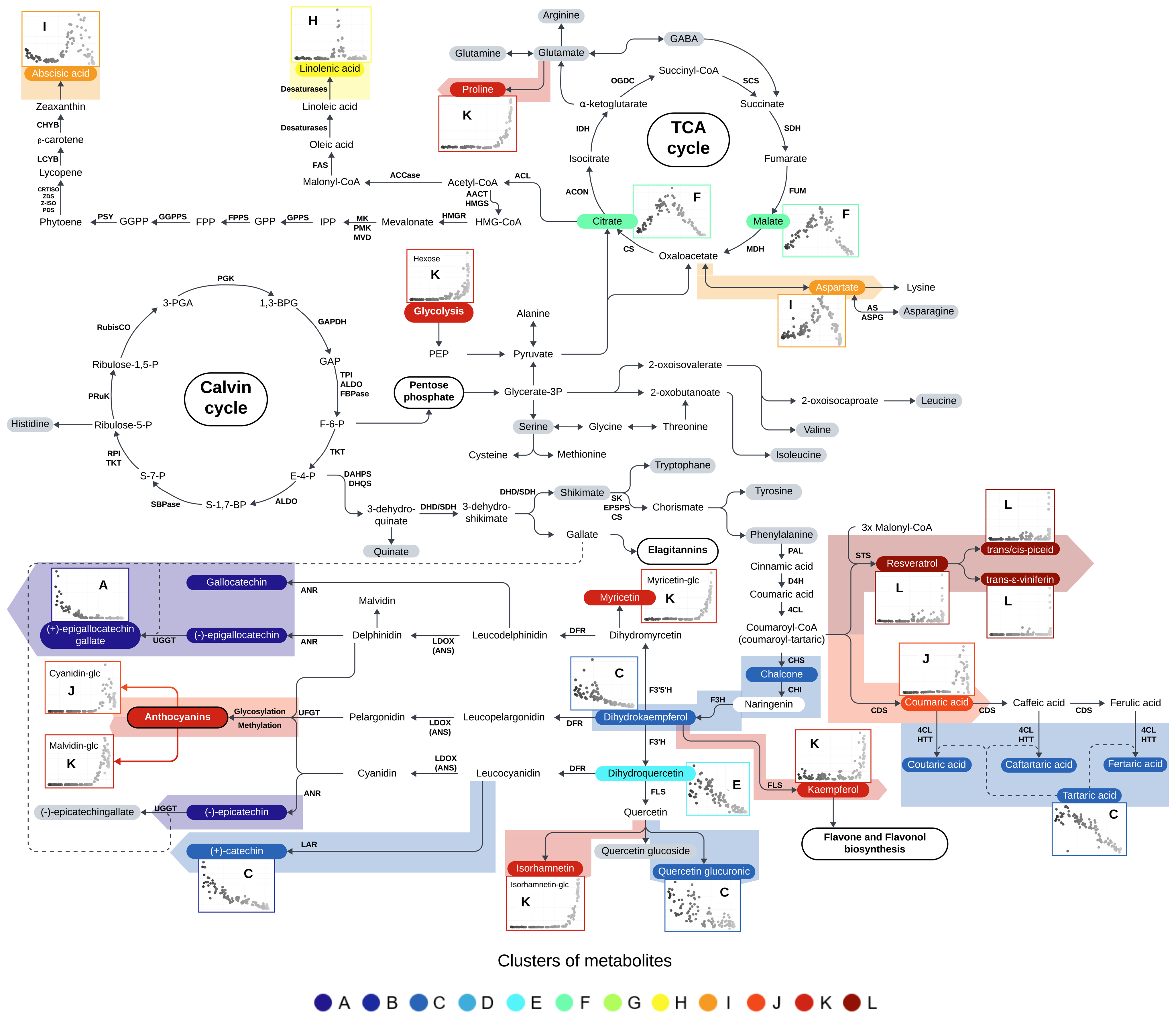
Biosynthesis pathways of the identified metabolites and their accumulation profile through the grapevine fruit development. Each point represents a metabolite, colored by its K-means cluster with its representative pattern over the metabolomic clock (see details in Fig. **S5**). Metabolites in gray correspond to molecular standards; most of these molecules have a correlation with the eigen-metabolite of their cluster of less than 0.8 (see dynamics in Fig. **S6**). Chalcone, epicatechin, gallocatechin, and epigallocatechin have been identified in their derivative, fragment, or polymerized forms. Due to the multiple profiles associated with each fragment/derivative/polymer, we have chosen not to display them in this figure (see details in Fig. **S5**). (KEGG Pathway database (https://www.genome.jp/kegg/pathway.html), Ali et al., 2010; Castellarin et al., 2007b; Dong and Lin, 2021; Clifford et al., 2017; Petrussa et al., 2013; Tang YuHan et al., 2018; Yang et al., 2020).

Catechins monomers are phenolic compounds produced via the flavonoid biosynthesis pathway, but their polymerisation in condensed tannins remains elusive (Yu et al., 2023). The conversion of phenylalanine into p-coumaroyl-CoA is considered as the first step in the phenylpropanoid pathway. The latter is then converted into flavonoids, including tannins, by chalcone synthase (CHS), or into stilbenes by stilbene synthase (STS) (Flamini et al., 2013; Rienth et al., 2021, Fig. 7). Profile A and C showed that the monomers and small proanthocyanidin (PA) polymers (EC, EGC, etc., from monomers to hexamers), rapidly decreased after anthesis. This decline coincided precisely with the period when higher molecular weight PAs began to accumulate, suggesting small polymers might be rate-limiting intermediaries (Kennedy et al., 2001, 2000; Ollé et al., 2011). Puzzlingly, profile A, encompassing many galloylated PA derivatives (Fig. **S3**), decreased before the non-galloylated ones in cluster C. Further work is needed to understand if the faster decrease of galloylated forms reflects (i) an increased in gallate/shikimate competition subsequent to an acceleration of the PA branch, (ii) an improved NADPH reducing power, or (iii) a sequential expression of specialized SDH/ isogenes (Tahara et al., 2020).

Linolenic acid, found in the H cluster announcing the onset of ripening at softening, is a precursor of jasmonic acid, a phytohormone involved in various developmental processes (Singh et al., 2022). It also serves as a precursor to diverse volatile compounds determinant for fruit aroma and defense mechanisms (Li et al., 2021; Rienth et al., 2021; Schwab et al., 2008). Its accumulation should mark increased membrane phospholipid turnover immediately before the onset of ripening, where its peroxidation takes place (Pilati et al., 2014).

The accumulation profiles of metabolites belonging to clusters J, K and L are consistent with their involvement in successive ripening-associated biological processes. During ripening, a second growth phase is set on due to a significant accumulation of water and solutes, mainly glucose and fructose, originating from sucrose translocated by phloem mass flow (Daviet et al., 2023). When the berries reach their maximum volume, phloem unloading in berries ceases, leading to a loss of water and volume known as the shriveling or over-ripening (Griesser et al., 2024; Rogiers et al., 2017; Savoi et al., 2021). The fruits then undergo constitutive water stress, which influences their metabolic composition.

### ABA as a key player of the ripening process

Cluster I started with the herbaceous plateau (L4) and peaked at the softening coloring transition (S7/R8). This cluster is marked by the accumulation of abscisic acid (ABA) in its free form, the level of which began to rise at the end of the herbaceous plateau, before peaking at the softening stage (Fig. **S5**). ABA is a phytohormone regulating plant growth and development. It plays multiple roles including tolerance to desiccation (Fujii and Zhu, 2009). ABA has been shown to promote color changes in non-climacteric fruits like strawberries (Batista-Silva et al., 2018; Jia et al., 2011; Jiang and Joyce, 2003). Analysis of grapevine berry development has highlighted ABA as an early indicator (if not a trigger) of veraison and berry coloration (Pilati et al., 2017), particularly by positively regulating the phenylpropanoid biosynthesis pathway (Chaves et al., 2010; Giribaldi et al., 2010; Kuhn et al., 2014; Lacampagne et al., 2010; Sun et al., 2010; Wheeler et al., 2009). This regulatory role of ABA is also evident in other non-climacteric fruits, such as strawberries (Jia et al., 2011), where ABA levels continue to increase from the beginning of ripening until the end of fruit development. In contrast, in grapevine berries, ABA peaks during the initial color change of the berries but then declines as the ripening phase progresses (Davies et al., 1997; Fortes et al., 2015; Pilati et al., 2017; Villalobos-González et al., 2016). Present single berry results largely refined these views, showing that ABA starts to accumulate before softening, acting as a trigger rather than a consequence of ripening, and maintains a high level throughout the entire phase of active phloem unloading, before shriveling (Fig. **S5**). If ABA belonged to cluster I, its glucose ester derivative (ABA-GE) accumulated later, and was found in cluster K. ABA-GE has been considered as a storage form of ABA or a final inactive product of its catabolism (Zeevaart, 1999). Hydrolysis of ABA-GE by specialized glucosidases has been shown to lead to a rapid increase in free ABA concentrations in response to osmotic stress in *Arabidopsis thaliana* (Xu et al., 2012). Moreover, the ABA catabolite phaseic acid was associated with cluster J (correlation of 0.69), suggesting that active ABA catabolism is responsible for the rapid decrease of free ABA at the very end of ripening (Fig. **S5**). Cluster J also contained a large number of sugar-derived ions, consistent with the accumulation profile of glucose and fructose as major *osmoticum* in ripening berries, starting at S6. To our knowledge, this is the first evidence that the sugar accumulation process precisely starts at the peak of ABA concentration (cluster I). ABA has been shown to activate the transcription of a cascade of enzyme and transporter genes playing key roles in sugar metabolism and accumulation during berry ripening (Bennett et al., 2023; Pan et al., 2005). Altogether, the rapid shift of the veraison-associated active ABA pool to inactive ABA forms such as ABA-GE and phaseic acid suggests that fine temporal tuning of ABA concentration is critical to a proper coordination of the ripening process.

### Flavonol and proline dynamics during ripening as indicators of constitutive stress

Noticeably, the sequential accumulation of anthocyanins and flavonols such as kaempferol, quercetin, isorhamnetin and myricetin derivatives, as well as malvidin and cyanidin derivatives in clusters J and K, reflects the decline of dihydroflavonols, such as dihydrokaempferol, in cluster C. This developmental shift in flavonol profiles follows the biosynthetic pathway and aligns with the expression profiles of the genes encoding dihydroflavonol-4-reductase and late genes of the anthocyanin regulated by MybA1 (Cutanda-Perez et al., 2009; Massonnet et al., 2017).

Cluster K shows a massive, but late accumulation of proline, already known to over-accumulate when both berry growth and net protein accumulation have ceased (Stines et al., 1999). The levels of delta-1-pyrroline-5-carboxylate synthetase mRNA and protein, a key regulatory enzyme in proline biosynthesis, do not seem to be affected by ripening (Deluc et al., 2009; Stines et al., 1999). Present results revealed that proline didn’t start to accumulate before late ripening (R8), and largely escaped the peak of free ABA, which in addition clearly vanished after phloem arrest, while proline accumulation persisted. This very poor coincidental timing between ABA and proline in single berries apparently contradicts the hypothesis that ABA would simply trigger proline biosynthesis in grapevines submitted to water deficit (Deluc et al., 2009). Finally, most metabolites associated to cluster L, which are characterized by a sharp accumulation at the very end of the ripening process, have been identified as stilbenes, including *trans*-resveratrol, *trans*-and *cis*-piceid and viniferins (Table **S2**). This cluster appeared therefore remarkably homogeneous, as all its annotated compounds, representing 83% of all ions, were derived from stilbenes (Table **S1**, Fig. **S3**). Stilbene biosynthesis has been associated with a wide array of biotic and abiotic stresses in grapevine (Chong et al., 2009). In grapes, stilbene metabolism has been shown to respond to water deficit, albeit with different responses depending on varieties (Deluc et al., 2011). In addition, transcriptomic analysis of berries subjected to post-harvest withering has shown a massive induction of the stilbene synthase gene family (Vannozzi et al., 2012). In the context of berry development, the present work shows that stilbenes can be considered as metabolic markers of the water stress associated to the cessation of phloem sap flow and wilting occuring at over-maturation stage.

### Dealing with complex accumulation profiles

K-means clustering, while useful for grouping data by similarity, may keep complex metabolic profiles hidden, resulting in misclassification of physiologically important metabolites. For example, by contrast with proline which also derives from glutamate, gamma-aminobutyric acid (GABA), the key intermediary of the GABA shunt in plants (Ansari et al., 2021), was found progressively accumulated during the green phase, followed by a sudden disappearance preceding softening, and a sharp increase at the end of maturation (Fig. **S6**). The recent identification of GABA as an inhibitor of the malate channel ALMT1 (Long et al., 2020), may provide a functional link between GABA abrupt decrease and the trigger of the typical reversal of malate accumulation during the green stage/ripening transition. Then, GABA being reversibly accumulated in mature berries submitted to O2 deprivation (Tesnière et al., 1994), its terminal accumulation is symptomatic of increased anaerobiosis in the fruit core at this stage (Xiao et al., 2018). Such a particular profile led to its misclassification in cluster E, albeit with a correlation of 0.62 to the cluster eigen-metabolite (Fig. **S6**). Similarly, tryptophan, a precursor of the phenylpropanoid pathway (Fig. 7), had a distinct profile with a moderated peak at veraison but an overall decreasing trend. This led to its placement in cluster D with a correlation of 0.64. To address this issue, increasing the number of clusters in K-means could allow for a finer classification of metabolites with specific trends. Furthermore, the use of more sophisticated grouping methodologies, for example by combining a Gaussian mixture model (Polanski et al., 2015) with the uniform or tree manifold approximation and projection (UMAP, TMAP) dimension reduction methods (Ebbels et al., 2023; Olivon et al., 2018), could allow better adaptation to these atypical profiles. This potential has already been partially explored in this study using t-SNE.

## Conclusion

This study provides a detailed analysis of grapevine berry development through an untargeted metabolomic profiling, identifying 12 distinct metabolite clusters that map the fruit’s phenological stages. Utilizing single berry sampling and multivariate analyses, we highlighted abrupt metabolic changes and the key roles of many metabolites. These results improve our understanding of the grape’s internal metabolic clock, offering the possibility of optimizing the harvesting of berries according to their internal developmental clock, thanks in particular to high-throughput and non-destructive phenotyping methods, opening the way to new applications in breeding. Additionally, this research paves the way for studying the physiology of non-climacteric fruits, allowing for more effective identification of key developmental regulators, such as the gene networks involved in the onset of ripening. This enhanced understanding could provide deeper insights into the molecular mechanisms underlying plant acclimatization to abiotic stresses.

## Acknowledgments

This work was supported by the Fondation Poupelain, the Agence Nationale de la Recherche (ANR, G2WAS project, ANR-19-CE20-0024), the Institut National de Recherche pour l’Agriculture, l’alimentation et l’Environnement (INRAe) Biologie et Amélioration des Plantes (BAP) department (Métab’EAU project) and the Région Occitanie (Métab’EAU project). Thanks to the INRAE UMR AGAP for helping with the sampling and to the INRAE UMR SVQV of Colmar for the metabolomics analysis and to Camille Rustenholz and Amandine Velt for the metabolomics pipeline.

## Author contributions

F.T. prepared the samples for LCMS analysis, established and implemented the multivariate analysis workflow, interpreted the results, and drafted the manuscript. S.S. monitored and collected the grapevine berry samples, prepared the samples, and contributed to the review and editing of the manuscript. L.T. provided the plant material and contributed to the review and editing of the manuscript. P.H. interpreted the results, and drafted the manuscript. R.B. supervised the metabolomic analyses, generated raw metabolomics data, identified the metabolites, interpreted the results, and drafted the manuscript. V.S. conceived and designed the study, coordinated and supervised the experiments, interpreted the results, and drafted the manuscript. C.R. monitored and collected the grapevine berry samples, conceived and designed the study, coordinated and supervised the experiments, interpreted the results, and drafted the manuscript. All authors have reviewed and approved the final version of the manuscript for publication.

## Data availability

Metabolomics data have been deposited to the EMBL-EBI MetaboLights database (DOI: 10.1093/nar/gkad1045, PMID:37971328) with the identifier MTBLS10572.

## References

Ali, K., Maltese, F., Choi, Y.H., Verpoorte, R., 2010. Metabolic constituents of grapevine and grape-derived products. Phytochem. Rev. 9, 357–378. 10.1007/s11101-009-9158-0

Ansari, M.I., Jalil, S.U., Ansari, S.A., Hasanuzzaman, M., 2021. GABA shunt: a key-player in mitigation of ROS during stress. Plant Growth Regul. 94, 131–149. 10.1007/s10725-021-00710-y

Batista-Silva, W., Nascimento, V.L., Medeiros, D.B., Nunes-Nesi, A., Ribeiro, D.M., Zsögön, A., Araújo, W.L., 2018. Modifications in Organic Acid Profiles During Fruit Development and Ripening: Correlation or Causation? Front. Plant Sci. 9. 10.3389/fpls.2018.01689

Bennett, J., Meiyalaghan, S., Nguyen, H.M., Boldingh, H., Cooney, J., Elborough, C., Araujo, L.D., Barrell, P., Lin-Wang, K., Plunkett, B.J., 2023. Exogenous abscisic acid and sugar induce a cascade of ripening events associated with anthocyanin accumulation in cultured Pinot Noir grape berries. Front. Plant Sci. 14, 1324675.

Bigard, A., Romieu, C., Sire, Y., Veyret, M., Ojeda, H., Torregrosa, L., 2019. The kinetics of grape ripening revisited through berry density sorting. OENO One 16.

Bonada, M., Sadras, V., Moran, M., Fuentes, S., 2013. Elevated temperature and water stress accelerate mesocarp cell death and shrivelling, and decouple sensory traits in Shiraz berries. Irrig. Sci. 31, 1317–1331. 10.1007/s00271-013-0407-z

Cabral, I.L., Teixeira, A., Ferrier, M., Lanoue, A., Valente, J., Rogerson, F.S., Alves, F., Carvalho, S.M.P., Gerós, H.V., Queiroz, J., 2023. Canopy management through crop forcing impacts grapevine cv. ‘Touriga Nacional’ performance, ripening and berry metabolomics profile. OENO One 57, 55–69. 10.20870/oeno-one.2023.57.1.7122

Castellarin, S.D., Di Gaspero, G., 2007. Transcriptional control of anthocyanin biosynthetic genes in extreme phenotypes for berry pigmentation of naturally occurring grapevines. BMC Plant Biol. 7, 46. 10.1186/1471-2229-7-46

Castellarin, S.D., Matthews, M.A., Di Gaspero, G., Gambetta, G.A., 2007a. Water deficits accelerate ripening and induce changes in gene expression regulating flavonoid biosynthesis in grape berries. Planta 227, 101–112.

Castellarin, S.D., Pfeiffer, A., Sivilotti, P., Degan, M., Peterlunger, E., Di Gaspero, G., 2007b. Transcriptional regulation of anthocyanin biosynthesis in ripening fruits of grapevine under seasonal water deficit. Plant Cell Environ. 30, 1381–1399.

Chaves, M.M., Zarrouk, O., Francisco, R., Costa, J.M., Santos, T., Regalado, A.P., Rodrigues, M.L., Lopes, C.M., 2010. Grapevine under deficit irrigation: hints from physiological and molecular data. Ann. Bot. 105, 661–676. 10.1093/aob/mcq030

Chong, J., Poutaraud, A., Hugueney, P., 2009. Metabolism and roles of stilbenes in plants. Plant Sci. 177, 143–155. 10.1016/j.plantsci.2009.05.012

Coombe, B. g., 1995. Growth Stages of the Grapevine: Adoption of a system for identifying grapevine growth stages. Aust. J. Grape Wine Res. 1, 104–110. 10.1111/j.1755-0238.1995.tb00086.x

Coombe, B.G., 1992. Research on Development and Ripening of the Grape Berry. Am. J. Enol. Vitic. 43, 101–110. 10.5344/ajev.1992.43.1.101

Coombe, B.G., 1976. The Development of Fleshy Fruits. Annu. Rev. Plant Physiol. 27, 207–228. 10.1146/annurev.pp.27.060176.001231

Coombe, B.G., Hale, C.R., 1973. The Hormone Content of Ripening Grape Berries and the Effects of Growth Substance Treatments. Plant Physiol. 51, 629–634. 10.1104/pp.51.4.629

CooMbe, B.G., McCarthy, M.G., 2000. Dynamics of grape berry growth and physiology of ripening. Aust. J. Grape Wine Res. 6, 131–135.

Cramer, G.R., Ghan, R., Schlauch, K.A., Tillett, R.L., Heymann, H., Ferrarini, A., Delledonne, M., Zenoni, S., Fasoli, M., Pezzotti, M., 2014. Transcriptomic analysis of the late stages of grapevine (Vitis vinifera cv. Cabernet Sauvignon) berry ripening reveals significant induction of ethylene signaling and flavor pathways in the skin. BMC Plant Biol. 14, 370. 10.1186/s12870-014-0370-8

Cutanda-Perez, M.-C., Ageorges, A., Gomez, C., Vialet, S., Terrier, N., Romieu, C., Torregrosa, L., 2009. Ectopic expression of VlmybA1 in grapevine activates a narrow set of genes involved in anthocyanin synthesis and transport. Plant Mol. Biol. 69, 633–648. 10.1007/s11103-008-9446-x

Dai, Z.W., Léon, C., Feil, R., Lunn, J.E., Delrot, S., Gomès, E., 2013. Metabolic profiling reveals coordinated switches in primary carbohydrate metabolism in grape berry (Vitis vinifera L.), a non-climacteric fleshy fruit. J. Exp. Bot. 64, 1345–1355. 10.1093/jxb/ers396

Davies, C., Robinson, S.P., Boss, P.K., 1997. Treatment of Crape Berries, a Nonclimacteric Fruit with a Synthetic Auxin, Retards Ripening and Alters the Expression of Developmentally Regulated Genes.

Daviet, B., Fournier, C., Cabrera-Bosquet, L., Simonneau, T., Cafier, M., Romieu, C., 2023. Ripening dynamics revisited: an automated method to track the development of asynchronous berries on time-lapse images. 10.1101/2023.07.12.548662

Del-Castillo-Alonso, M.-Á., Monforte, L., Tomás-Las-Heras, R., Ranieri, A., Castagna, A., Martínez-Abaigar, J., Núñez-Olivera, E., 2021. Secondary metabolites and related genes in Vitis vinifera L. cv. Tempranillo grapes as influenced by ultraviolet radiation and berry development. Physiol. Plant. 173, 709–724. 10.1111/ppl.13483

Deluc, L.G., Decendit, A., Papastamoulis, Y., Mérillon, J.-M., Cushman, J.C., Cramer, G.R., 2011. Water Deficit Increases Stilbene Metabolism in Cabernet Sauvignon Berries. J. Agric. Food Chem. 59, 289–297. 10.1021/jf1024888

Deluc, L.G., Quilici, D.R., Decendit, A., Grimplet, J., Wheatley, M.D., Schlauch, K.A., Mérillon, J.-M., Cushman, J.C., Cramer, G.R., 2009. Water deficit alters differentially metabolic pathways affecting important flavor and quality traits in grape berries of Cabernet Sauvignon and Chardonnay. BMC Genomics 10, 212. 10.1186/1471-2164-10-212

Dong, N.-Q., Lin, H.-X., 2021. Contribution of phenylpropanoid metabolism to plant development and plant–environment interactions. J. Integr. Plant Biol. 63, 180–209. 10.1111/jipb.13054

Duan, S., Wu, Y., Fu, R., Wang, L., Chen, Y., Xu, W., Zhang, C., Ma, C., Shi, J., Wang, S., 2019. Comparative Metabolic Profiling of Grape Skin Tissue along Grapevine Berry Developmental Stages Reveals Systematic Influences of Root Restriction on Skin Metabolome. Int. J. Mol. Sci. 20, 534. 10.3390/ijms20030534

Ebbels, T.M.D., van der Hooft, J.J.J., Chatelaine, H., Broeckling, C., Zamboni, N., Hassoun, S., Mathé, E.A., 2023. Recent advances in mass spectrometry-based computational metabolomics. Curr. Opin. Chem. Biol. 74, 102288. 10.1016/j.cbpa.2023.102288

Fasoli, M., Dal Santo, S., Zenoni, S., Tornielli, G.B., Farina, L., Zamboni, A., Porceddu, A., Venturini, L., Bicego, M., Murino, V., Ferrarini, A., Delledonne, M., Pezzotti, M., 2012. The Grapevine Expression Atlas Reveals a Deep Transcriptome Shift Driving the Entire Plant into a Maturation Program. Plant Cell 24, 3489–3505. 10.1105/tpc.112.100230

Fasoli, M., Richter, C.L., Zenoni, S., Bertini, E., Vitulo, N., Dal Santo, S., Dokoozlian, N., Pezzotti, M., Tornielli, G.B., 2018. Timing and Order of the Molecular Events Marking the Onset of Berry Ripening in Grapevine. Plant Physiol. 178, 1187–1206. 10.1104/pp.18.00559

Flamini, R., Mattivi, F., Rosso, M.D., Arapitsas, P., Bavaresco, L., 2013. Advanced Knowledge of Three Important Classes of Grape Phenolics: Anthocyanins, Stilbenes and Flavonols. Int. J. Mol. Sci. 14, 19651–19669. 10.3390/ijms141019651

Fortes, A.M., Teixeira, R.T., Agudelo-Romero, P., 2015. Complex Interplay of Hormonal Signals during Grape Berry Ripening. Molecules 20, 9326–9343. 10.3390/molecules20059326

Fujii, H., Zhu, J.-K., 2009. Arabidopsis mutant deficient in 3 abscisic acid-activated protein kinases reveals critical roles in growth, reproduction, and stress. Proc. Natl. Acad. Sci. 106, 8380–8385. 10.1073/pnas.0903144106

Gillaspy, G., Ben-David, H., Gruissem, W., 1993. Fruits: A Developmental Perspective. Plant Cell 5, 1439–1451.

Giovannoni, J.J., 2004. Genetic Regulation of Fruit Development and Ripening. Plant Cell 16, S170–S180. 10.1105/tpc.019158

Giribaldi, M., Gény, L., Delrot, S., Schubert, A., 2010. Proteomic analysis of the effects of ABA treatments on ripening Vitis vinifera berries. J. Exp. Bot. 61, 2447–2458. 10.1093/jxb/erq079

Goes da Silva, F., Iandolino, A., Al-Kayal, F., Bohlmann, M.C., Cushman, M.A., Lim, H., Ergul, A., Figueroa, R., Kabuloglu, E.K., Osborne, C., Rowe, J., Tattersall, E., Leslie, A., Xu, J., Baek, J., Cramer, G.R., Cushman, J.C., Cook, D.R., 2005. Characterizing the Grape Transcriptome. Analysis of Expressed Sequence Tags from Multiple Vitis Species and Development of a Compendium of Gene Expression during Berry Development. Plant Physiol. 139, 574–597. 10.1104/pp.105.065748

Govender, P., Sivakumar, V., 2020. Application of *k*-means and hierarchical clustering techniques for analysis of air pollution: A review (1980–2019). Atmospheric Pollut. Res. 11, 40–56. 10.1016/j.apr.2019.09.009

Griesser, M., Savoi, S., Bondada, B., Forneck, A., Keller, M., 2024. Berry shrivel in grapevine: a review considering multiple approaches. J. Exp. Bot. 75, 2196–2213. 10.1093/jxb/erae001

Hebra, T., Elie, N., Poyer, S., Van Elslande, E., Touboul, D., Eparvier, V., 2021. Dereplication, Annotation, and Characterization of 74 Potential Antimicrobial Metabolites from Penicillium Sclerotiorum Using t-SNE Molecular Networks. Metabolites 11, 444. 10.3390/metabo11070444

Jia, H.-F., Chai, Y.-M., Li, C.-L., Lu, D., Luo, J.-J., Qin, L., Shen, Y.-Y., 2011. Abscisic Acid Plays an Important Role in the Regulation of Strawberry Fruit Ripening. Plant Physiol. 157, 188–199. 10.1104/pp.111.177311

Jiang, Y., Joyce, D.C., 2003. ABA effects on ethylene production, PAL activity, anthocyanin and phenolic contents of strawberry fruit. Plant Growth Regul. 39, 171–174. 10.1023/A:1022539901044

Kennard, R.W., Stone, L.A., 1969. Computer Aided Design of Experiments. Technometrics 11, 137–148. 10.1080/00401706.1969.10490666

Kennedy, J.A., Hayasaka, Y., Vidal, S., Waters, E.J., Jones, G.P., 2001. Composition of Grape Skin Proanthocyanidins at Different Stages of Berry Development. J. Agric. Food Chem. 49, 5348–5355. 10.1021/jf010758h

Kennedy, J.A., Troup, G.J., Pilbrow, J.R., Hutton, D.R., Hewitt, D., Hunter, C.R., Ristic, R., Iland, P.G., Jones, G.P., 2000. Development of seed polyphenols in berries from Vitis vinifera L. cv. Shiraz. Aust. J. Grape Wine Res. 6, 244–254. 10.1111/j.1755-0238.2000.tb00185.x

Kuhn, N., Guan, L., Dai, Z.W., Wu, B.-H., Lauvergeat, V., Gomès, E., Li, S.-H., Godoy, F., Arce-Johnson, P., Delrot, S., 2014. Berry ripening: recently heard through the grapevine. J. Exp. Bot. 65, 4543–4559. 10.1093/jxb/ert395

Lacampagne, S., Gagné, S., Gény, L., 2010. Involvement of Abscisic Acid in Controlling the Proanthocyanidin Biosynthesis Pathway in Grape Skin: New Elements Regarding the Regulation of Tannin Composition and Leucoanthocyanidin Reductase (LAR) and Anthocyanidin Reductase (ANR) Activities and Expression. J. Plant Growth Regul. 29, 81–90. 10.1007/s00344-009-9115-6

Leng, F., Duan, S., Song, S., Zhao, L., Xu, W., Zhang, C., Ma, C., Wang, L., Wang, S., 2021. Comparative Metabolic Profiling of Grape Pulp during the Growth Process Reveals Systematic Influences under Root Restriction. Metabolites 11, 377. 10.3390/metabo11060377

Li, S., Chen, K., Grierson, D., 2021. Molecular and Hormonal Mechanisms Regulating Fleshy Fruit Ripening. Cells 10, 1136. 10.3390/cells10051136

Long, Y., Tyerman, S.D., Gilliham, M., 2020. Cytosolic GABA inhibits anion transport by wheat ALMT1. New Phytol. 225, 671–678. 10.1111/nph.16238

Lorenz, D. h., Eichhorn, K. w., Bleiholder, H., Klose, R., Meier, U., Weber, E., 1995. Growth Stages of the Grapevine: Phenological growth stages of the grapevine (Vitis vinifera L. ssp. vinifera)—Codes and descriptions according to the extended BBCH scale†. Aust. J. Grape Wine Res. 1, 100–103. 10.1111/j.1755-0238.1995.tb00085.x

Lund, S.T., Bohlmann, J., 2006. The Molecular Basis for Wine Grape Quality-A Volatile Subject. Science 311, 804–805. 10.1126/science.1118962

Massonnet, M., Fasoli, M., Tornielli, G.B., Altieri, M., Sandri, M., Zuccolotto, P., Paci, P., Gardiman, M., Zenoni, S., Pezzotti, M., 2017. Ripening Transcriptomic Program in Red and White Grapevine Varieties Correlates with Berry Skin Anthocyanin Accumulation. Plant Physiol. 174, 2376–2396. 10.1104/pp.17.00311

Matthews, M., Cheng, G., Weinbaum, S., 1987. Changes in water potential and dermal extensibility during grape berry development. J. Am. Soc. Hortic. Sci. 112, 314–319.

N. Clifford, M., B. Jaganath, I., A. Ludwig, I., Crozier, A., 2017. Chlorogenic acids and the acyl-quinic acids: discovery, biosynthesis, bioavailability and bioactivity. Nat. Prod. Rep. 34, 1391–1421. 10.1039/C7NP00030H

Nicolas, S., Bois, B., Billet, K., Romanet, R., Bahut, F., Uhl, J., Schmitt-Kopplin, P., Gougeon, R.D., 2024. High-Resolution Mass Spectrometry-Based Metabolomics for Increased Grape Juice Metabolite Coverage. Foods 13, 54. 10.3390/foods13010054

Ojeda, H., Deloire, A., Carbonneau, A., Ageorges, A., Romieu, C., 1999. Berry development of grapevines: relations between the growth of berries and their DNA content indicate cell multiplication and enlargement. VITIS-J. Grapevine Res. 38, 145.

Olivon, F., Elie, N., Grelier, G., Roussi, F., Litaudon, M., Touboul, D., 2018. MetGem Software for the Generation of Molecular Networks Based on the t-SNE Algorithm. Anal. Chem. 90, 13900–13908. 10.1021/acs.analchem.8b03099

Ollé, D., Guiraud, J. l., Souquet, J. m., Terrier, N., Ageorges, A., Cheynier, V., Verries, C., 2011. Effect of pre- and post-veraison water deficit on proanthocyanidin and anthocyanin accumulation during Shiraz berry development. Aust. J. Grape Wine Res. 17, 90–100. 10.1111/j.1755-0238.2010.00121.x

Pan, Q.-H., Li, M.-J., Peng, C.-C., Zhang, N., Zou, X., Zou, K.-Q., Wang, X.-L., Yu, X.-C., Wang, X.-F., Zhang, D.-P., 2005. Abscisic acid activates acid invertases in developing grape berry. Physiol. Plant. 125, 157–170. 10.1111/j.1399-3054.2005.00552.x

Petrussa, E., Braidot, E., Zancani, M., Peresson, C., Bertolini, A., Patui, S., Vianello, A., 2013. Plant Flavonoids—Biosynthesis, Transport and Involvement in Stress Responses. Int. J. Mol. Sci. 14, 14950–14973. 10.3390/ijms140714950

Pilati, S., Bagagli, G., Sonego, P., Moretto, M., Brazzale, D., Castorina, G., Simoni, L., Tonelli, C., Guella, G., Engelen, K., Galbiati, M., Moser, C., 2017. Abscisic Acid Is a Major Regulator of Grape Berry Ripening Onset: New Insights into ABA Signaling Network. Front. Plant Sci. 8. 10.3389/fpls.2017.01093

Pilati, S., Brazzale, D., Guella, G., Milli, A., Ruberti, C., Biasioli, F., Zottini, M., Moser, C., 2014. The onset of grapevine berry ripening is characterized by ROS accumulation and lipoxygenase-mediated membrane peroxidation in the skin. BMC Plant Biol. 14, 87. 10.1186/1471-2229-14-87

Polanski, A., Marczyk, M., Pietrowska, M., Widlak, P., Polanska, J., 2015. Signal Partitioning Algorithm for Highly Efficient Gaussian Mixture Modeling in Mass Spectrometry. PLOS ONE 10, e0134256. 10.1371/journal.pone.0134256

Rienth, M., Vigneron, N., Darriet, P., Sweetman, C., Burbidge, C., Bonghi, C., Walker, R.P., Famiani, F., Castellarin, S.D., 2021. Grape Berry Secondary Metabolites and Their Modulation by Abiotic Factors in a Climate Change Scenario–A Review. Front. Plant Sci. 12. 10.3389/fpls.2021.643258

Rodrigues, M., Forestan, C., Ravazzolo, L., Hugueney, P., Baltenweck, R., Rasori, A., Cardillo, V., Carraro, P., Malagoli, M., Brizzolara, S., Quaggiotti, S., Porro, D., Meggio, F., Bonghi, C., Battista, F., Ruperti, B., 2023. Metabolic and Molecular Rearrangements of Sauvignon Blanc (Vitis vinifera L.) Berries in Response to Foliar Applications of Specific Dry Yeast. Plants 12, 3423. 10.3390/plants12193423

Rogiers, S.Y., Coetzee, Z.A., Walker, R.R., Deloire, A., Tyerman, S.D., 2017. Potassium in the Grape (Vitis vinifera L.) Berry: Transport and Function. Front. Plant Sci. 8. 10.3389/fpls.2017.01629

Rousseeuw, P.J., 1987. Silhouettes: A graphical aid to the interpretation and validation of cluster analysis. J. Comput. Appl. Math. 20, 53–65. 10.1016/0377-0427(87)90125-7

Savoi, S., Torregrosa, L., Romieu, C., 2023. Single berry development – a new phenotyping and transcriptomics paradigm. VITIS - J. Grapevine Res. 62, 49–55. 10.5073/vitis.2023.62.special-issue.49-55

Savoi, S., Torregrosa, L., Romieu, C., 2021. Transcripts switched off at the stop of phloem unloading highlight the energy efficiency of sugar import in the ripening V. vinifera fruit. 10.1101/2021.01.19.427234

Schwab, W., Davidovich-Rikanati, R., Lewinsohn, E., 2008. Biosynthesis of plant-derived flavor compounds. Plant J. 54, 712–732. 10.1111/j.1365-313X.2008.03446.x

Schymanski, E.L., Jeon, J., Gulde, R., Fenner, K., Ruff, M., Singer, H.P., Hollender, J., 2014. Identifying Small Molecules via High Resolution Mass Spectrometry: Communicating Confidence. Environ. Sci. Technol. 48, 2097–2098. 10.1021/es5002105

Seymour, G.B., Østergaard, L., Chapman, N.H., Knapp, S., Martin, C., 2013. Fruit Development and Ripening. Annu. Rev. Plant Biol. 64, 219–241. 10.1146/annurev-arplant-050312-120057

Shahood, R., Torregrosa, L., Savoi, S., Romieu, C., 2020. First quantitative assessment of growth, sugar accumulation and malate breakdown in a single ripening berry. Oeno One 54, 1077–1092.

Singh, P., Arif, Y., Miszczuk, E., Bajguz, A., Hayat, S., 2022. Specific Roles of Lipoxygenases in Development and Responses to Stress in Plants. Plants 11, 979. 10.3390/plants11070979

Sorkun, M.C., Mullaj, D., Koelman, J.V.A., Er, S., 2022. ChemPlot, a Python library for chemical space visualization.

Stines, A.P., Naylor, D.J., Høj, P.B., van Heeswijck, R., 1999. Proline Accumulation in Developing Grapevine Fruit Occurs Independently of Changes in the Levels of Δ1-Pyrroline-5-Carboxylate Synthetase mRNA or Protein1. Plant Physiol. 120, 923. 10.1104/pp.120.3.923

Sun, L., Zhang, M., Ren, J., Qi, J., Zhang, G., Leng, P., 2010. Reciprocity between abscisic acid and ethylene at the onset of berry ripening and after harvest. BMC Plant Biol. 10, 257. 10.1186/1471-2229-10-257

Tahara, K., Nishiguchi, M., Funke, E., Miyazawa, S.-I., Miyama, T., Milkowski, C., 2020. Dehydroquinate dehydratase/shikimate dehydrogenases involved in gallate biosynthesis of the aluminum-tolerant tree species Eucalyptus camaldulensis. Planta 253, 3. 10.1007/s00425-020-03516-w

Tang YuHan, T.Y., Jiang Yao, J.Y., Meng JiaSong, M.J., Tao Jun, T.J., 2018. A brief review of physiological roles, plant resources, synthesis, purification and oxidative stability of alpha-linolenic acid.

Terrier, N., Romieu, C., 2001. Grape Berry Acidity, in: Roubelakis-Angelakis, K.A. (Ed.), Molecular Biology & Biotechnology of the Grapevine. Springer Netherlands, Dordrecht, pp. 35–57. 10.1007/978-94-017-2308-4_2

Tesnière, C., Romieu, C., Dugelay, I., Nicol, M.Z., Flanzy, C., Robin, J.P., 1994. Partial recovery of grape energy metabolism upon aeration following anaerobic stress. J. Exp. Bot. 45, 145–151. 10.1093/jxb/45.1.145

Thomas, T.R., Shackel, K.A., Matthews, M.A., 2008. Mesocarp cell turgor in Vitis vinifera L. berries throughout development and its relation to firmness, growth, and the onset of ripening. Planta 228, 1067–1076. 10.1007/s00425-008-0808-z

Tornielli, G.B., Sandri, M., Fasoli, M., Amato, A., Pezzotti, M., Zuccolotto, P., Zenoni, S., 2023. A molecular phenology scale of grape berry development. Hortic. Res. 10, uhad048. 10.1093/hr/uhad048

Vannozzi, A., Dry, I.B., Fasoli, M., Zenoni, S., Lucchin, M., 2012. Genome-wide analysis of the grapevine stilbene synthase multigenic family: genomic organization and expression profiles upon biotic and abiotic stresses. BMC Plant Biol. 12, 130. 10.1186/1471-2229-12-130

Villalobos-González, L., Peña-Neira, A., Ibáñez, F., Pastenes, C., 2016. Long-term effects of abscisic acid (ABA) on the grape berry phenylpropanoid pathway: Gene expression and metabolite content. Plant Physiol. Biochem. 105, 213–223. 10.1016/j.plaphy.2016.04.012

Wheeler, S., Loveys, B., Ford, C., Davies, C., 2009. The relationship between the expression of abscisic acid biosynthesis genes, accumulation of abscisic acid and the promotion of Vitis vinifera L. berry ripening by abscisic acid. Aust. J. Grape Wine Res. 15, 195–204. 10.1111/j.1755-0238.2008.00045.x

Xiao, Z., Rogiers, S.Y., Sadras, V.O., Tyerman, S.D., 2018. Hypoxia in grape berries: the role of seed respiration and lenticels on the berry pedicel and the possible link to cell death. J. Exp. Bot. 69, 2071–2083. 10.1093/jxb/ery039

Xu, Z.-Y., Lee, K.H., Dong, T., Jeong, J.C., Jin, J.B., Kanno, Y., Kim, D.H., Kim, S.Y., Seo, M., Bressan, R.A., Yun, D.-J., Hwang, I., 2012. A Vacuolar β-Glucosidase Homolog That Possesses Glucose-Conjugated Abscisic Acid Hydrolyzing Activity Plays an Important Role in Osmotic Stress Responses in Arabidopsis. Plant Cell 24, 2184–2199. 10.1105/tpc.112.095935

Yang, B., He, S., Liu, Y., Liu, B., Ju, Y., Kang, D., Sun, X., Fang, Y., 2020. Transcriptomics integrated with metabolomics reveals the effect of regulated deficit irrigation on anthocyanin biosynthesis in Cabernet Sauvignon grape berries. Food Chem. 314, 126170. 10.1016/j.foodchem.2020.126170

Yu, K., Song, Y., Lin, J., Dixon, R.A., 2023. The complexities of proanthocyanidin biosynthesis and its regulation in plants. Plant Commun. 4, 100498. 10.1016/j.xplc.2022.100498

Zamboni, A., Di Carli, M., Guzzo, F., Stocchero, M., Zenoni, S., Ferrarini, A., Tononi, P., Toffali, K., Desiderio, A., Lilley, K.S., Pè, M.E., Benvenuto, E., Delledonne, M., Pezzotti, M., 2010. Identification of Putative Stage-Specific Grapevine Berry Biomarkers and Omics Data Integration into Networks. Plant Physiol. 154, 1439–1459. 10.1104/pp.110.160275

Zeevaart, J.A.D., 1999. Chapter 8 - Abscisic acid metabolism and its regulation, in: Hooykaas, P.J.J., Hall, M.A., Libbenga, K.R. (Eds.), New Comprehensive Biochemistry, Biochemistry and Molecular Biology of Plant Hormones. Elsevier, pp. 189–207. 10.1016/S0167-7306(08)60488-3

