## Supplementary material for "The single-berry metabolomic clock paradigm reveals new stages and metabolic switches during grapevine berry development"

### Supporting Information

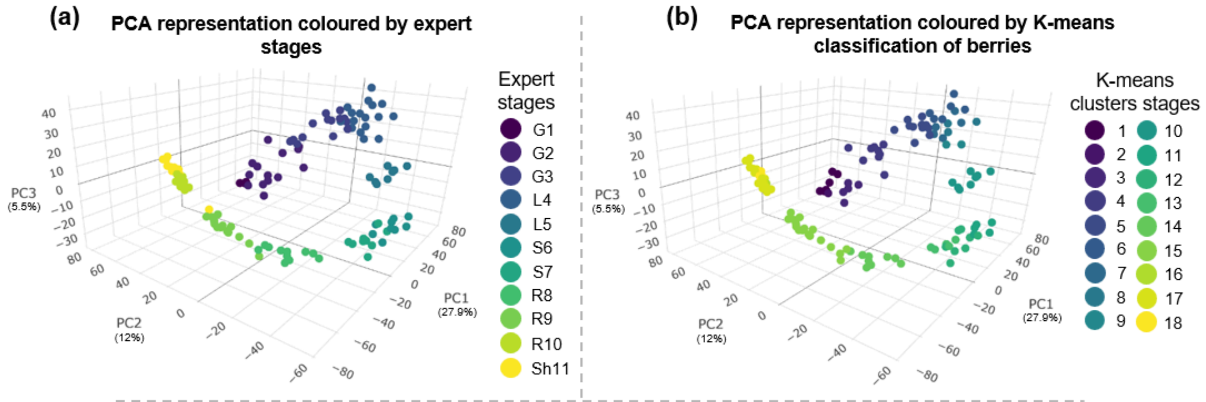


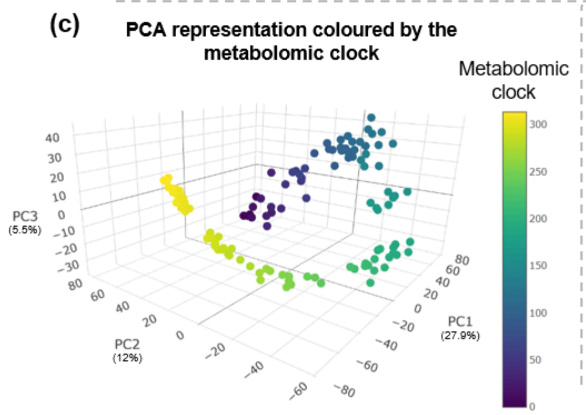


**Figure S1. Comparative analysis of berry developmental stages: expert classification (A), K-means method (B) and metabolomic clock (C).** G = green, L = lag, S = softening, R = ripening, Sh = shriveling. Each point represents a single berry. The PCA plots illustrate the separation and distribution of berries within the multi-dimensional space defined by the first three principal components of the dataset.

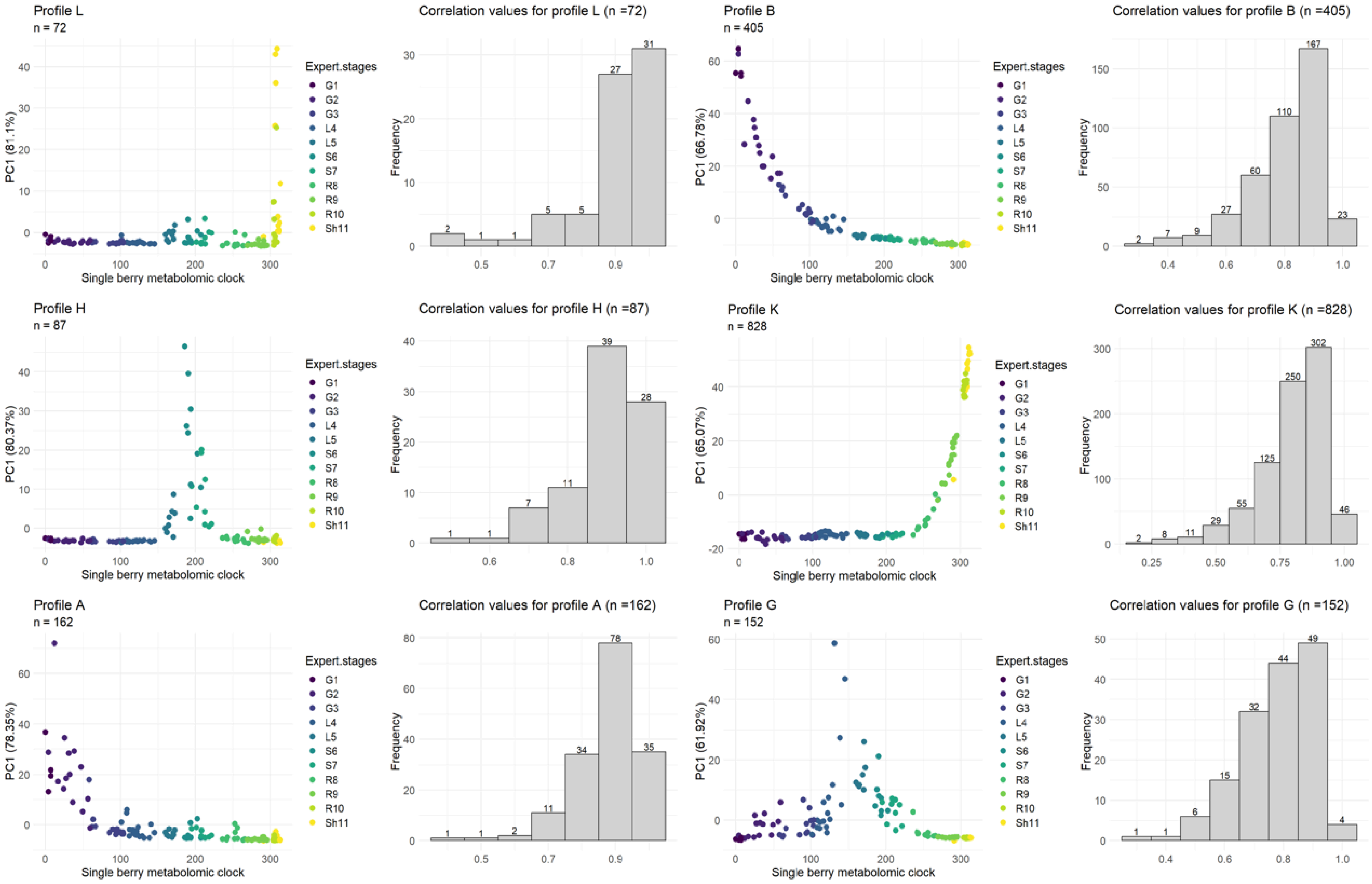


**Figure S2. Correlation of all the metabolites in the cluster to the representative curve (from the best to the worst).**

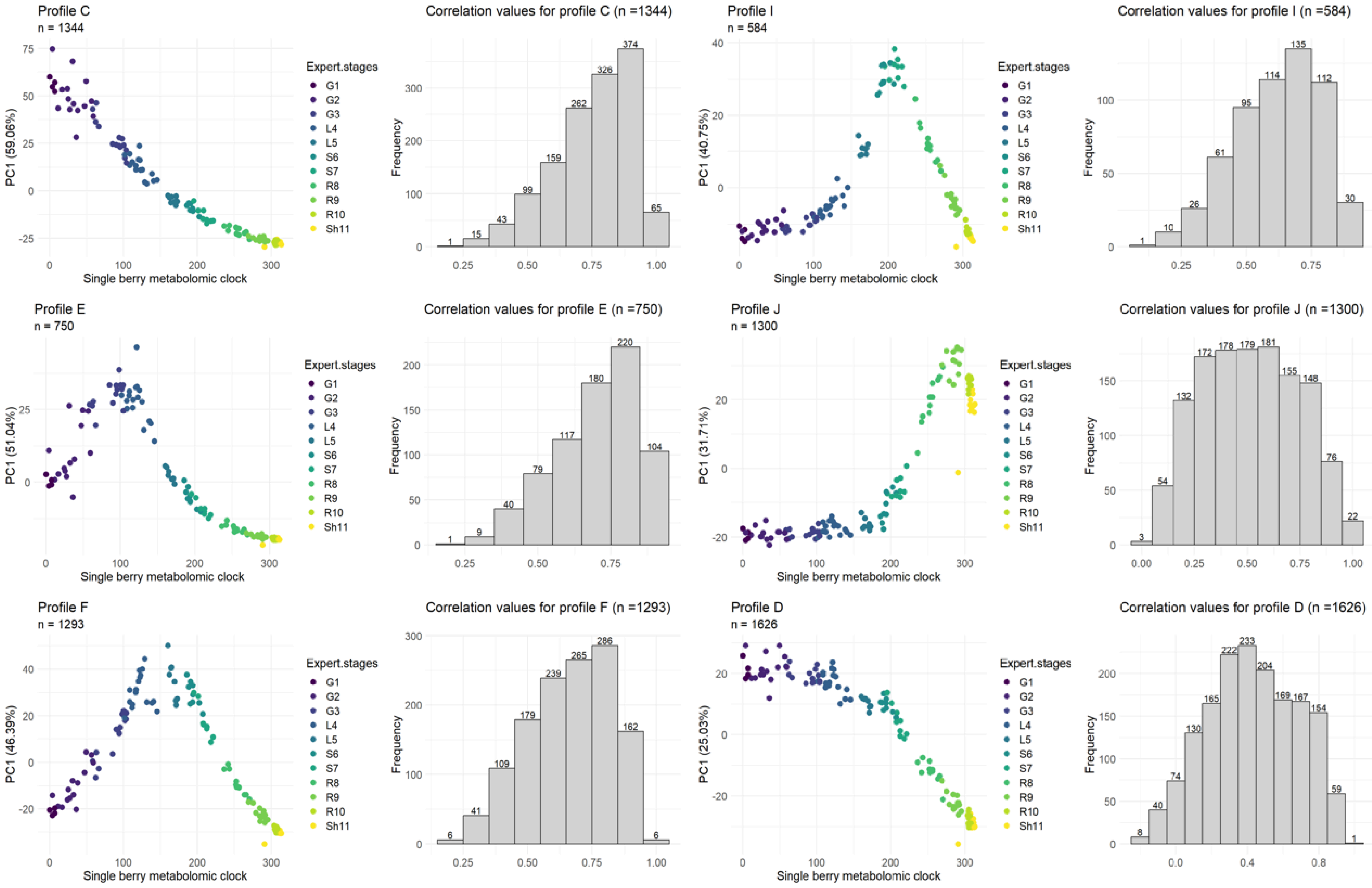


**Figure S2 *bis*. Correlation of all the metabolites in the cluster to the representative curve (from the best to the worst).**

**Table S1. Annotation of selected ions in clusters of interest.** Ions whose correlation with the eigen-metabolite of each cluster was greater than or equal to 0.8 were selected in positive and negative mode and subjected to expert annotation. The proportion of annotated ions in each cluster is indicated, as well as the main chemical families of the corresponding metabolites.

| Clusters | Number of ions in positive mode | Number of ions in negative mode | Total of ions | Main chemical families of the corresponding metabolites | Annotated ions | % annotated ions |
|---|---|---|---|---|---|---|
| A | 32 | 105 | 137 | Galloyl derivatives of condensed tannins | 53 | 39 |
| B | 88 | 166 | 254 | / | 0 | 0 |
| C | 305 | 298 | 603 | Condensed tannins, flavonoids | 152 | 25 |
| D | 99 | 46 | 145 | Condensed tannins | 12 | 8 |
| E | 128 | 85 | 213 | Condensed tannins, flavonoids | 14 | 7 |
| F | 235 | 78 | 313 | Organic acids | 24 | 8 |
| G | 54 | 21 | 75 | / | 0 | 0 |
| H | 68 | 8 | 76 | Lipids | 10 | 13 |
| I | 48 | 26 | 74 | Amino acids, sugars, hormone | 16 | 22 |
| J | 108 | 51 | 159 | Sugars, anthocyanin | 56 | 35 |
| K | 126 | 351 | 477 | Proline, sugars, flavonoids, inorganic acids, anthocyanins | 96 | 20 |
| L | 19 | 41 | 60 | Stilbenes | 50 | 83 |
| **Total** | **1310** | **1276** | **2586** | | **483** | **19** |

**Table S2. Annotation of the most correlated metabolites to their cluster's representative curve.** Metabolites were identified based on expert analysis of MS and MS/MS spectra. Ion mass to charge ratios (*m/z* in u) and retention times (RT, in min) are indicated. The "Corr" column is the correlation value of the ion dynamic with the eigen-metabolite dynamic of its cluster. Abbreviations: Glc: glucoside, GlcA: glucuronide. Links of the different online databases: PubChem (https://pubchem.ncbi.nlm.nih.gov/), ChemSpider (http://www.chemspider.com/), mzCloud (https://www.mzcloud.org/), FoodDB (https://bmjopen.bmj.com/content/9/6/e026652), KNApSAcK Family (http://www.knapsackfamily.com/KNApSAcK_Family/) and MassBank (https://massbank.eu/MassBank/).

| Name | Family | Formula | RT (min) | Positive mode | | | Negative mode | | | Validation | | ICL | Cluster | literature |
|---|---|---|---|---|---|---|---|---|---|---|---|---|---|---|
| | | | | Cor | m/z (u) | | Cor | m/z (u) | | Standard | Online database | | | |
| Epiafzelechin - 3 - O - gallate | Tannin gallate ester | C22H18O9 | 1,92 | 0,97 | 427,10258 | [M+H]+ | 0,97 | 425,08747 | [M-H]- | no | PubChem; ChemSpider | 3 | A | |
| Epicatechin tetramer | Condensed tannin | C60H50O24 | 1,94 | | | | 0,97 | 577,63097 | [M-2H]2- | no | mzCloud; PubChem; ChemSpider | 2 | A | Jouin et al., 2022 |
| Epicatechin hexamere digallate | Tannin gallate ester | C104H84O44 | 2,01 | | | | 0,94 | 1017,72282 | [M-2H]2- | no | | 3 | A | |
| Procyanidin B2 | Condensed tannin | C30H28O12 | 2,11 | | | | 0,85 | 579,15056 | [M-H]- | yes | mzCloud; PubChem; ChemSpider | 1 | A | Di Lecce et al., 2014 |
| Epicatechin trimer 3 - gallate | Tannin gallate ester | C52H42O22 | 2,51 | 0,84 | 1019,22413 | [M+H]+ | | | | no | FoodDB; KNApSAcK Family; PubChem | 2 | A | Rockenbach et al., 2012 |
| Epigallocatechine gallate | Tannin gallate ester | C22H18 O11 | 2,55 | | | | 0,88 | 457,07738 | [M-H]- | yes | mzCloud; PubChem; ChemSpider | 1 | A | Di Lecce et al., 2014 |
| Procyanidin B 3 - gallate | Tannin gallate ester | C37H30O16 | 2,69 | 0,99 | 731,16044 | [M+H]+ | | | | no | PubChem; ChemSpider | 2 | A | Rockenbach et al., 2012 |
| Epicatechin trimer digallate | Tannin gallate ester | C59H46O26 | 3,12 | 0,96 | 1171,23657 | [M+H]+ | | | | no | mzCloud; PubChem; ChemSpider | 3 | A | Rockenbach et al., 2012 |
| Epicatechin hexamere digallate | Tannin gallate ester | C104H86O44 | 3,19 | 0,92 | 1018,21568 | [M+2H]2+ | | | | no | | 4 | A | |
| Epicatechin trimer 3 - gallate | Tannin gallate ester | C52H42O22 | 3,23 | | | | 0,93 | 1017,20808 | [M-H]- | no | FoodDB; KNApSAcK Family; PubChem | 3 | A | Rockenbach et al., 2012 |

| | | | | | | | | | | | | | | |
|---|---|---|---|---|---|---|---|---|---|---|---|---|---|---|
| Procyanidin type B 3,3' - di - O - gallate | Tannin gallate ester | C44H35O20 | 3,39 | 0,95 | 883,17181 | [M+H]+ | | | | no | mzCloud; PubChem; ChemSpider | 3 | A | Rockenbach et al., 2012 |
| Aromadendrin gallate | Tannin gallate ester | C22H16O10 | 3,41 | 0,94 | 441,08189 | [M+H]+ | | | | no | PubChem | 4 | A | |
| Gallocatechine gallate | Tannin gallate ester | C22H18O11 | 3,48 | | | | 0,82 | 457,07726 | [M-H]- | yes | mzCloud; PubChem | 3 | A | Rockenbach et al., 2012; Di Lecce et al., 2014 |
| Epicatechin hexamere digallate | Tannin gallate ester | C104H88O44 | 3,63 | 0,98 | 1018,2147 | [M+2H]2+ | | | | no | PubChem | 4 | A | |
| Epicatechin trimer 3 - gallate | Tannin gallate ester | C52H42O22 | 3,64 | 0,99 | 1019,22421 | [M+H]+ | 0,99 | 1017,20805 | [M-H]- | no | FoodDB; KNApSAcK Family; PubChem | 3 | A | Rockenbach et al., 2012 |
| Procyanidin B 3 - gallate | Tannin gallate ester | C37H30O16 | 3,87 | 0,95 | 731,16064 | [M+H]+ | | | | no | PubChem; ChemSpider | 3 | A | Rockenbach et al., 2012 |
| Epicatechin trimer 3 - gallate | Tannin gallate ester | C52H42O22 | 3,96 | 0,88 | 1019,22405 | [M+H]+ | | | | no | PubChem; ChemSpider | 3 | A | Rockenbach et al., 2012 |
| Epicatechin gallate | Tannin gallate ester | C22H18O10 | 3,98 | | | | 0,86 | 441,08209 | [M-H]- | yes | mzCloud; PubChem; ChemSpider | 1 | A | Rockenbach et al., 2012 |
| Epicatechin trimer 3 - gallate | Tannin gallate ester | C52H42O22 | 3,99 | | | | 0,94 | 1017,20705 | [M-H]- | no | PubChem; ChemSpider | 3 | A | Rockenbach et al., 2012 |
| Aromadendrin gallate | Tannin gallate ester | C22H16O10 | 4,30 | 0,93 | 441,08181 | [M+H]+ | | | | no | PubChem | 4 | A | de Oliveira et al., 2019 |
| Epicatechin hexamere digallate | Tannin gallate ester | C104H88O44 | 4,57 | 0,96 | 1019,22414 | [M+2H]2+ | | | | no | FoodDB; KNApSAcK Family; PubChem | 4 | A | |
| Epicatechin trimer 3 - gallate | Tannin gallate ester | C52H42O22 | 4,66 | | | | 0,93 | 1017,20698 | [M-H]- | no | FoodDB; KNApSAcK Family; PubChem | 3 | A | Rockenbach et al., 2012 |
| Procyanidin B 3 - gallate | Tannin gallate ester | C37H30O16 | 4,66 | 0,98 | 731,16068 | [M+H]+ | | | | no | PubChem; ChemSpider | 3 | A | Rockenbach et al., 2012 |
| Aromadendrin gallate | Tannin gallate ester | C22H16O10 | 4,74 | | | | 0,81 | 439,0667 | [M-H]- | no | | 4 | A | de Oliveira et al., 2019 |
| Epicatechin tetramer 3 - gallate | Tannin gallate ester | C67H54O28 | 4,85 | | | | 0,97 | 652,13247 | [M-2H]2- | no | | 3 | A | Rockenbach et al., 2012; Frost et al., 2018 |
| Epicatechin trimer digallate | Tannin gallate ester | C59H46O26 | 4,95 | | | | 0,89 | 584,10618 | [M-2H]2- | no | FoodDB; KNApSAcK Family; PubChem | 3 | A | Rockenbach et al., 2012 |

| | | | | | | | | | | | | | | |
|---|---|---|---|---|---|---|---|---|---|---|---|---|---|---|
| Procyanidin B 3,3' - di - O - gallate | Tannin gallate ester | C44H34O20 | 5,42 | 0,94 | 883,17217 | [M+H]+ | 0,95 | 881,15604 | [M-H]- | no | mzCloud; PubChem; ChemSpider | 3 | A | Rockenbach et al., 2012 |
| Procyanidin type C | Condensed tannin | C45H38O18 | 1,25 | 0,90 | 867,21373 | [M+H]+ | | | | no | mzCloud; PubChem; ChemSpider | 2 | C | Goufo et al., 2020 |
| Procyanidin type C | Condensed tannin | C45H38O18 | 1,58 | 0,88 | 867,21377 | [M+H]+ | | | | yes | mzCloud; PubChem; ChemSpider | 2 | C | Goufo et al., 2020 |
| Catechin - catechin - gallocatechin trimer | Condensed tannin | C45 H38 O19 | 1,70 | 0,87 | 883,2085 | [M+H]+ | | | | no | mzCloud; PubChem; ChemSpider | 2 | C | Frost et al., 2018 |
| Procyanidin type A (dimer) | Condensed tannin | C30 H24O12 | 1,91 | 0,85 | 577,13435 | [M+H]+ | | | | yes | mzCloud; PubChem; ChemSpider | 1 | C | J. Int. Sci. Vigne Vin, 2001, 35, n°1, 51-56 |
| Procyanidin B2 | Condensed tannin | C30 H26 O12 | 1,93 | 0,81 | 579,14975 | [M+H]+ | | | | yes | mzCloud; PubChem; ChemSpider | 1 | C | Goufo et al., 2020 |
| Procyanidin type C | Condensed tannin | C45H38O18 | 1,93 | 0,92 | 867,21351 | [M+H]+ | 0,92 | 865,19788 | [M-H]- | no | mzCloud; PubChem; ChemSpider | 2 | C | Goufo et al., 2020 |
| Catechin - gallocatechin dimer | Condensed tannin | C30H24O13 | 1,94 | | | | 0,92 | 591,11441 | [M-H]- | no | mzCloud; PubChem; ChemSpider | 2 | C | Goufo et al., 2020 |
| Procyanidin type C (trimer) | Condensed tannin | C46 H40 O20 | 1,99 | | | | 0,85 | 911,20329 | [M-H]- | no | mzCloud; PubChem; ChemSpider | 2 | C | Goufo et al., 2020 |
| Procyanidin type A (dimer) | Condensed tannin | C30H24O12 | 2,11 | 0,83 | 577,13422 | [M+H]+ | | | | no | mzCloud; PubChem; ChemSpider | 2 | C | Goufo et al., 2020 |
| Catechin | Condensed tannin | C15H14O6 | 2,12 | | | | 0,92 | 289,07127 | [M-H]- | yes | mzCloud; PubChem; ChemSpider | 1 | C | Goufo et al., 2020 |
| Procyanidin type C (trimer) | Condensed tannin | C46H40O20 | 2,14 | 0,87 | 913,21951 | [M+H]+ | 0,88 | 911,20311 | [M-H]- | no | | 2 | C | Goufo et al., 2020 |
| Taxifolin fragment (3) | Flavonoid | C15H12O7 | 2,26 | 0,83 | 305,06561 | [M+H]+ | 0,89 | 303,05077 | [M-H]- | no | | 4 | C | Goufo et al., 2020 |
| Caftaric acid | Phenolic acid | C13H12O9 | 2,34 | 0,92 | 313,0555 | [M+H]+ | 0,96 | 311,04041 | [M-H]- | yes | mzCloud; PubChem; ChemSpider | 1 | C | Lago-Vanzela et al., 2011 |
| Procyanidin C2 | Condensed tannin | C45H38O18 | 2,33 | 0,84 | 867,2137 | [M+H]+ | 0,81 | 865,19711 | [M-H]- | yes | mzCloud; PubChem; ChemSpider | 1 | C | Goufo et al., 2020 |
| Procyanidin type C (trimer) | Condensed tannin | C46H40O19 | 2,60 | 0,95 | 897,22441 | [M+H]+ | 0,94 | 895,20861 | [M-H]- | no | | 2 | C | Goufo et al., 2020 |

| | | | | | | | | | | | | | | |
|---|---|---|---|---|---|---|---|---|---|---|---|---|---|---|
| Procyanidin B1 | Condensed tannin | C30H26O12 | 2,65 | 0,91 | 579,14982 | [M+H]+ | 0,86 | 577,13482 | [M-H]- | yes | mzCloud; PubChem; ChemSpider | 1 | C | Goufo et al., 2020 |
| Coutaric acid | Phenolic acid | C13H11O8 | 3,03 | 0,97 | 319,0425 | [M+H]+ | 0,98 | 295,04538 | [M-H]- | yes | mzCloud; PubChem; ChemSpider | 1 | C | Lago-Vanzela et al., 2011 |
| Chalcone - Glc derivative (1) | Flavonoid | C21H22O10 | 3,33 | 0,84 | 435,1287 | [M+H]+ | | | | no | | 4 | C | |
| Kaempferol - Glc derivative | Flavonoid | C21H20O11 | 3,69 | 0,92 | 449,1081 | [M+H]+ | 0,94 | 447,09289 | [M-H]- | no | | 4 | C | Kedrina-Okutan et al., 2018 |
| Dihydrokaempferol - Glc | Flavonoid | C21H22H11 | 3,74 | 0,82 | 451,1241 | [M+H]+ | 0,82 | 449,10868 | [M-H]- | yes | mzCloud; PubChem; ChemSpider | 1 | C | Kedrina-Okutan et al., 2018 |
| Fertaric acid | Phenolic acid | C14H14O9 | 3,85 | | | | 0,95 | 325,05624 | [M-H]- | yes | mzCloud; PubChem; ChemSpider | 1 | C | Lago-Vanzela et al., 2011 |
| Quercetin - GlcA | Flavonoid | C21H18O13 | 4,08 | 0,81 | 479,08187 | [M+H]+ | 0,82 | 477,06672 | [M-H]- | yes | mzCloud; PubChem; ChemSpider | 2 | C | Benbouguerra et al., 2021 |
| Dihydrochalcone - Glc derivative | Flavonoid | C21H24O10 | 4,30 | 0,87 | 437,14482 | [M+H]+ | 0,90 | 435,12968 | [M-H]- | no | | 4 | C | |
| Chalcone - Glc derivative (2) | Flavonoid | C21H22O10 | 4,31 | 0,94 | 435,12884 | [M+H]+ | 0,95 | 433,11364 | [M-H]- | no | | 4 | C | |
| Chalcone - Glc derivative (3) | Flavonoid | C21H22O10 | 4,97 | 0,95 | 435,12874 | | | | | no | | 4 | C | |
| Tartaric acid | Organic acid | C4H6O6 | 1,41 | | | | 0,88 | 149,00891 | [M-H]- | yes | mzCloud; PubChem; ChemSpider | 1 | D | *Preiner et al., 2013* |
| Procyanidin type B | Condensed tannin | C30H26O12 | 1,72 | 0,94 | 579.14964 | [M+H]+ | 0,85 | 577,13486 | [M-H]- | no | mzCloud; PubChem; ChemSpider | 2 | D | Goufo et al., 2020 |
| Procyanidin type A (dimer) | Condensed tannin | C30H24O12 | 1,41 | 0,88 | 577.13413 | [M+H]+ | | | | no | mzCloud; PubChem; ChemSpider | 2 | D | Goufo et al., 2020; J. Int. Sci. Vigne Vin, 2001, 35, n°1, 51-56 |
| Taxifolin fragment (1) | Flavonoid | C15H12O7 | 1,76 | 0,92 | 305,06563 | [M+H]+ | | | | no | | 4 | E | Goufo et al., 2020 |
| Taxifolin fragment (2) | Flavonoid | C15H12O7 | 1,41 | 0,85 | 305,06559 | [M+H]+ | | | | no | | 4 | E | Goufo et al., 2020 |
| Prodelphinidin type B | Condensed tannin | C30H26O14 | 1,25 | 0,83 | 611.13965 | [M+H]+ | | | | no | mzCloud; PubChem; ChemSpider | 2 | E | Bindon et al., 2014 |
| Catechin - gallocatechin dimer | Condensed tannin | C30H26O13 | 1,26 | 0,92 | 595.1448 | [M+H]+ | | | | no | mzCloud; PubChem; ChemSpider | 2 | E | Bindon et al., 2014 |
| Catechin - gallocatechin dimer | Condensed tannin | C30H26O13 | 1,59 | 0,91 | 595.14467 | [M+H]+ | | | | no | mzCloud; PubChem; ChemSpider | 2 | E | Bindon et al., 2014 |

| | | | | | | | | | | | | | | |
|---|---|---|---|---|---|---|---|---|---|---|---|---|---|---|
| Citric acid | organic acid | C6H8O7 | 1,44 | 0,92 | 193,03429 | [M+H]+ | 0,91 | 191,01951 | [M-H]- | yes | mzCloud; PubChem; ChemSpider | 1 | F | *Preiner et al., 2013* |
| Malic acid | organic acid | C4H6O5 | 1,6 | 0,89 | 135,02878 | [M+H]+ | 0,96 | 133,01409 | [M-H]- | yes | mzCloud; PubChem; ChemSpider | 1 | F | *Preiner et al., 2013* |
| Linolenic acid | Lipid | C18H30O2 | 10,24 | 0,98 | 279,23221 | [M+H]+ | | | | yes | PubChem | 1 | H | Iglesias et al., 1991 |
| C18:3 acid | Lipid | C18H30O2 | 10,45 | 0,94 | 279,23206 | [M+H]+ | | | | no | PubChem | 2 | H | Iglesias et al., 1991 |
| Dihomo - $\gamma$ - linolenic acid | Lipid | C20H34O2 | 11,01 | 0,94 | 307,26332 | [M+H]+ | | | | no | PubChem | 3 | H | |
| Dihomo - $\gamma$ - linolenic acid | Lipid | C20H34O2 | 11,85 | 0,86 | 324,28982 | [M+NH4]+ | | | | no | PubChem | 3 | H | |
| diterpenol (geranyllinalool) | Terpene | C20H34O | 10,86 | 0,97 | 291,2683 | [M+H]+ | | | | yes | | 2 | H | |
| Aspartic acid | Amino acid | C4H7O4N | 1,08 | 0,90 | 134,04479 | [M+H]+ | | | | yes | mzCloud; PubChem; ChemSpider | 1 | I | Bouzas-Cid et al., 2015 |
| Hexose | Sugar | C6H12O6 | 1,16 | | | | 0,82 | 179,0559 | [M-H]- | yes | mzCloud; PubChem; ChemSpider | 2 | I | |
| Polyol | Sugar | C22H38O21 | 1,16 | | | | 0,83 | 637,18274 | [M-H]- | no | mzCloud; PubChem; ChemSpider | 2 | I | |
| Hexose tetramer | Sugar | C24H43O22 | 1,11 | | | | 0,86 | 683,22433 | [M-H]- | no | | 2 | J | |
| Hexose dimer | Sugar | C12H22O11 | 1,11 | 0,82 | 365,10556 | | 0,82 | 341,10823 | [M-H]- | yes | | 2 | J | |
| Cyanidin - Glc | Anthocyanin | C21H20O11 | 1,59 | 0,82 | 449,10807 | [M+H]+ | | | | yes | mzCloud; PubChem; ChemSpider | 1 | J | Benbouguerra et al., 2021 |
| Coumaric - Glc - Ester | Phenolic acid | C15H18O8 | 6,45 | | | | 0,80 | 325,09268 | [M-H]- | no | | 2 | J | Di Lecce et al., 2014 |
| Dihydro - isorhamnetin derivative | Flavonoid | C16H14O7 | 2,38 | 0,83 | 319,0813 | [M+H]+ | | | | no | | 3 | J | De Rosso et al., 2014 |
| Dihydrosyringetin | Flavonoid | C17H16O8 | 5,34 | | | | 0,88 | 347,07683 | [M-H]- | no | | 3 | J | Narduzzi et al., 2015 |
| Glucosaminyl - myo - inositol | Sugar | C12H23O10N | 0,89 | 0,86 | 342,13956 | [M+H]+ | | | | no | ChemSpider | 3 | K | |
| N - methylethanolamine phosphate | Lipid | C3H10NO4P | 0,89 | 0,94 | 156,04218 | [M+H]+ | | | | no | | 4 | K | |
| Potassium | Inorganic ion | CHO2K2 | 0,92 | 0,90 | 122,92446 | [M+AF]+ | | | | yes | | 1 | K | |
| Phosphoric acid | Inorganic ion | H3PO4 | 1,53 | | | | 0,91 | 96.96949 | [M-H]- | yes | | 1 | K | |
| Proline | Amino Acid | C5H9NO2 | 1,08 | 0,97 | 116,07051 | [M+H]+ | 0,85 | 114,05596 | [M-H]- | yes | mzCloud; PubChem; ChemSpider | 1 | K | |

| | | | | | | | | | | | | | | |
|---|---|---|---|---|---|---|---|---|---|---|---|---|---|---|
| Fructosyl proline | Amino Acid | C11H19O7N | 1,09 | 0,93 | 278,12345 | [M+H]+ | | | | no | | 3 | K | Hashiba 1978 |
| Hexose | Sugar | C6 H12 O6 | 1,15 | 0,84 | 219,02656 | [M+K]+ | | | | yes | | 1 | K | |
| Hexose trimer | Sugar | C18H32O16 | 1,20 | 0,90 | 543,13233 | [M+K]+ | | | | yes | | 1 | K | |
| Hexose dimer | Sugar | C12H22O11 | 1,30 | 0,92 | 381,0794 | [M+K]+ | | | | yes | | 1 | K | |
| Malvidin - Glc | Anthocyanin | C23H25O12 | 1,25 | 0,82 | 493,13411 | [M+H]+ | | | | yes | mzCloud; PubChem; MassBank | 1 | K | Benbouguerra et al., 2021 |
| Glucose - 6 - phosphate derivative | Sugar | C6H14O10P | 1,53 | | | | 0,97 | 277,0327 | [M-H]- | no | | 2 | K | Diakou et al., 1997 |
| Peonidin - Acetyl - Glc | Anthocyanin | C24H25O12 | 2,09 | 0,85 | 505,13406 | [M+H]+ | | | | no | mzCloud; PubChem; Phytohub | 2 | K | Benbouguerra et al., 2021 |
| Malvidin - Acetyl - Glc | Anthocyanin | C25H27O13 | 2,15 | 0,93 | 535,14445 | [M+H]+ | | | | no | | 2 | K | Benbouguerra et al., 2021 |
| Myricetin - Glc | Flavonoid | C21H20O13 | 2,73 | 0,98 | 481,09775 | [M+H]+ | 0,98 | 479,08255 | [M-H]- | no | | 2 | K | de Oliveira et al., 2019 |
| Abscisic acid - Glc | Hormone | C21H30O9 | 3,90 | 0,92 | 449,17842 | [M+H]+ | | | | no | | 2 | K | Gagné et al., 2006; Priest et al., 2005 |
| Malvidin - Coumaroyl - Glc | Anthocyanin | C32H31O14 | 3,95 | 0,97 | 639,17038 | [M+H]+ | | | | no | | 2 | K | Benbouguerra et al., 2021 |
| Kaempferol - hexoside | Flavonoid | C21H20O11 | 4,22 | 0,85 | 449,1081 | [M+H]+ | | | | no | | 3 | K | Liang et al., 2012 |
| Kaempferol - Glc | Flavonoid | C21H20O11 | 4,49 | 0,87 | 449,10806 | [M+H]+ | 0,87 | 447,09279 | [M-H]- | yes | | 1 | K | Benbouguerra et al., 2021 |
| Isorhamnetin - Glc | Flavonoid | C22H22O12 | 4,60 | 0,98 | 479,11845 | [M+H]+ | 0,97 | 477,1032 | [M-H]- | yes | | 1 | K | de Oliveira et al., 2019 |
| Isorhamnetin - hexoside | Flavonoid | C22H22O12 | 4,99 | | | | 0,87 | 477,10362 | [M-H]- | no | | 3 | K | de Oliveira et al., 2019 |
| Isorhamnetin - GlcA | Flavonoid | C22H20O13 | 5,00 | 0,89 | 493,09791 | [M+H]+ | 0,87 | 491,08287 | [M-H]- | no | | 3 | K | de Oliveira et al., 2019 |
| Isorhamnetin - hexoside | Flavonoid | C22H22O12 | 5,38 | | | | 0,83 | 477,10369 | [M-H]- | no | | 3 | K | de Oliveira et al., 2019 |
| Isorhamnetin - pentoside | Flavonoid | C21H20O11 | 5,44 | 0,85 | 449,10813 | [M+H]+ | 0,87 | 447,09304 | [M-H]- | no | | 3 | K | Mattivi et al., 2006 |
| Astringin (Piceatannol-Glc) | Stilbene | C20H22O9 | 2,54 | | | | 0,87 | 405,1196 | [M-H]- | no | mzCloud; PubChem; ChemSpider | 2 | L | Guerrero et al., 2020 |
| *trans* - Resveratrol - GlcA | Stilbene | C20H20O9 | 2,55 | | | | 0,96 | 403,10303 | [M-H]- | no | | 3 | L | Muzzio et al., 2012 |
| Restrytisol B | Stilbene | C28H24O7 | 3,56 | | | | 0,94 | 471,14461 | [M-H]- | no | | 2 | L | Cichewicz et al., 2000 |
| *trans* - Piceid | Stilbene | C20H22O8 | 3,71 | 0,98 | 229,08594 | [M+H]+ | 0,97 | 389,12369 | [M-H]- | yes | mzCloud; PubChem; ChemSpider | 1 | L | Chong et al., 2009 |

| | | | | | | | | | | | | | | |
|---|---|---|---|---|---|---|---|---|---|---|---|---|---|---|
| *cis* - Resveratrol GlcA | Stilbene | $C_{20}H_{20}O_9$ | 3,78 | | | | 0,95 | 403,103 | [M-H]- | no | | 3 | L | Muzzio et al., 2012 |
| Astringin (piceatannol - Glc) | Stilbene | $C_{20}H_{22}O_9$ | 4,00 | | | | 0,91 | 405,1188 | [M-H]- | no | mzCloud; PubChem; ChemSpider | 2 | L | Chong et al., 2009 |
| Resveratrol - GlcA | Stilbene | $C_{20}H_{20}O_9$ | 4,01 | | | | 0,92 | 403,10298 | [M-H]- | no | | 2 | L | |
| Resveratrol - Rut | Stilbene | $C_{26}H_{32}O_{12}$ | 4,31 | | | | 0,96 | 535,18182 | [M-H]- | no | ChemSpider | 2 | L | Aja et al., 2019 |
| Resveratroloside | Stilbene | $C_{20}H_{22}O_8$ | 4,61 | | | | 0,92 | 389,12384 | [M-H]- | no | mzCloud, ChemSpider | 2 | L | Aja et al., 2019 |
| Dehydroresveratrol - Glc (1) | Stilbene | $C_{20}H_{20}O_8$ | 5,02 | 0,90 | 227,07033 | [M+H]+ | 0,92 | 387,10804 | [M-H]- | no | | 4 | L | |
| *cis* - Piceid | Stilbene | $C_{20}H_{22}O_8$ | 5,11 | 0,99 | 413,12099 | [M+Na]+ | 0,99 | 389,12357 | [M-H]- | yes | | 1 | L | Chong et al., 2009 |
| δ - viniferin or pallidol like | Stilbene | $C_{28}H_{22}O_6$ | 5,80 | 0,95 | 455,149 | [M+H]+ | | | | no | mzCloud, ChemSpider | 2 | L | He et al., 2009; Pezet et al., 2003 |
| Dehydroresveratrol - Glc (2) | Stilbene | $C_{20}H_{20}O_8$ | 5,83 | | | | 0,95 | 387,10806 | [M-H]- | no | | 4 | L | |
| Gnetin F like | Stilbene | $C_{28}H_{24}O_6$ | 5,85 | | | | 0,93 | 455,09573 | [M-H]- | no | mzCloud, ChemSpider | 4 | L | |
| δ - viniferin or pallidol like | Stilbene | $C_{28}H_{22}O_6$ | 5,86 | | | | 0,94 | 453,134 | [M-H]- | no | mzCloud, ChemSpider | 2 | L | He et al., 2009; Pezet et al., 2003 |
| Deoxyrhapontigenin Rut | Stilbene | $C_{27}H_{34}O_{12}$ | 6,31 | | | | 0,85 | 549,1975 | [M-H]- | no | | 3 | L | Hoferer et al., 2023 |
| Viniferin - Glc | Stilbene | $C_{34}H_{32}O_{11}$ | 6,41 | 0,87 | 617,20162 | [M+H]+ | | | | no | mzCloud, ChemSpider | 3 | L | Aja et al., 2019 |
| *trans* - Resveratrol | Stilbene | $C_{14}H_{12}O_3$ | 6,54 | 0,92 | 229,0861 | [M+H]+ | | | | yes | mzCloud, ChemSpider | 1 | L | Chong et al., 2009 |

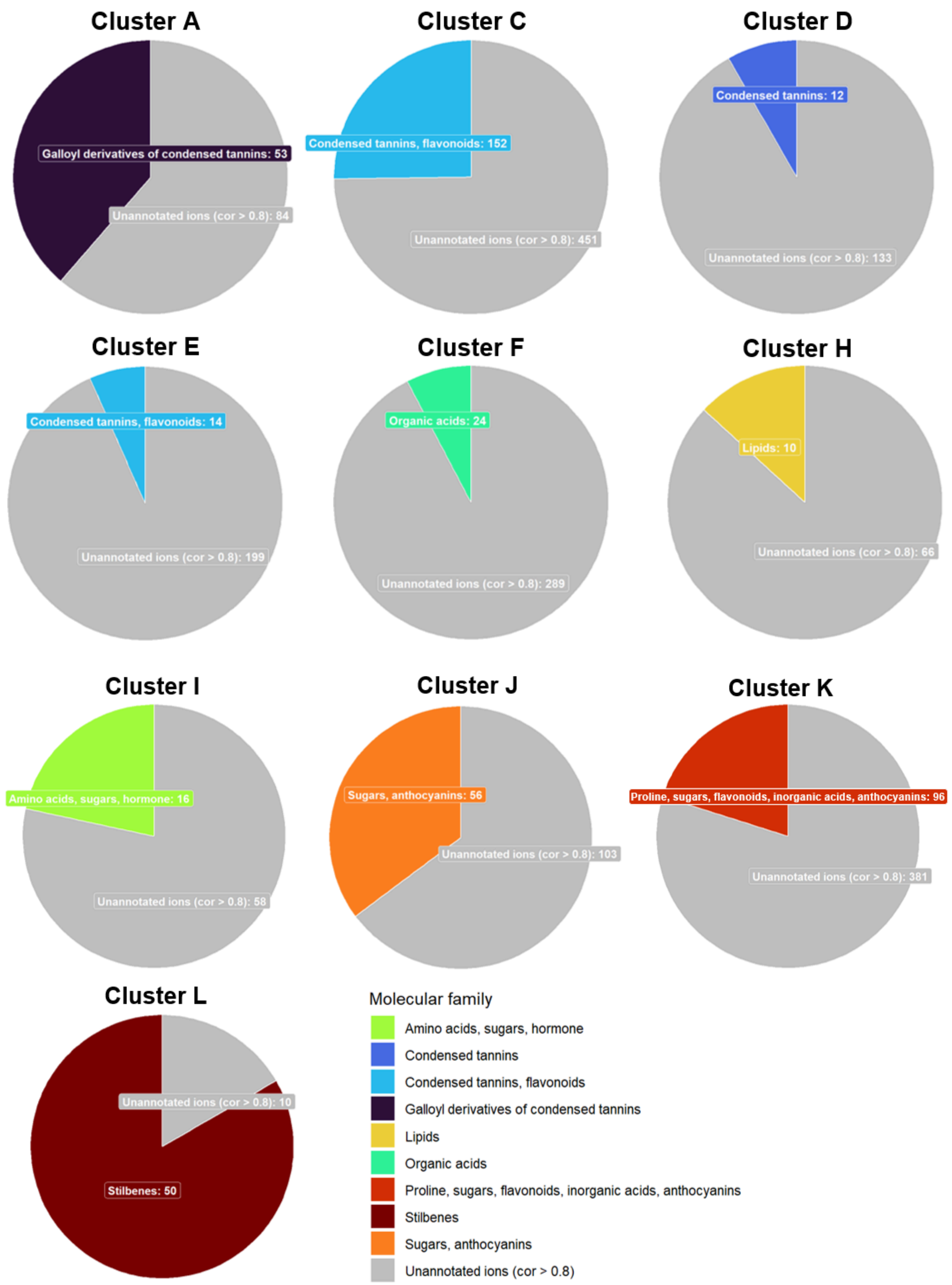


**Figure S3. Proportion of annotated ions in the selected clusters (cor > 0.8).** The chemical families, to which the identified ions belong, are indicated.

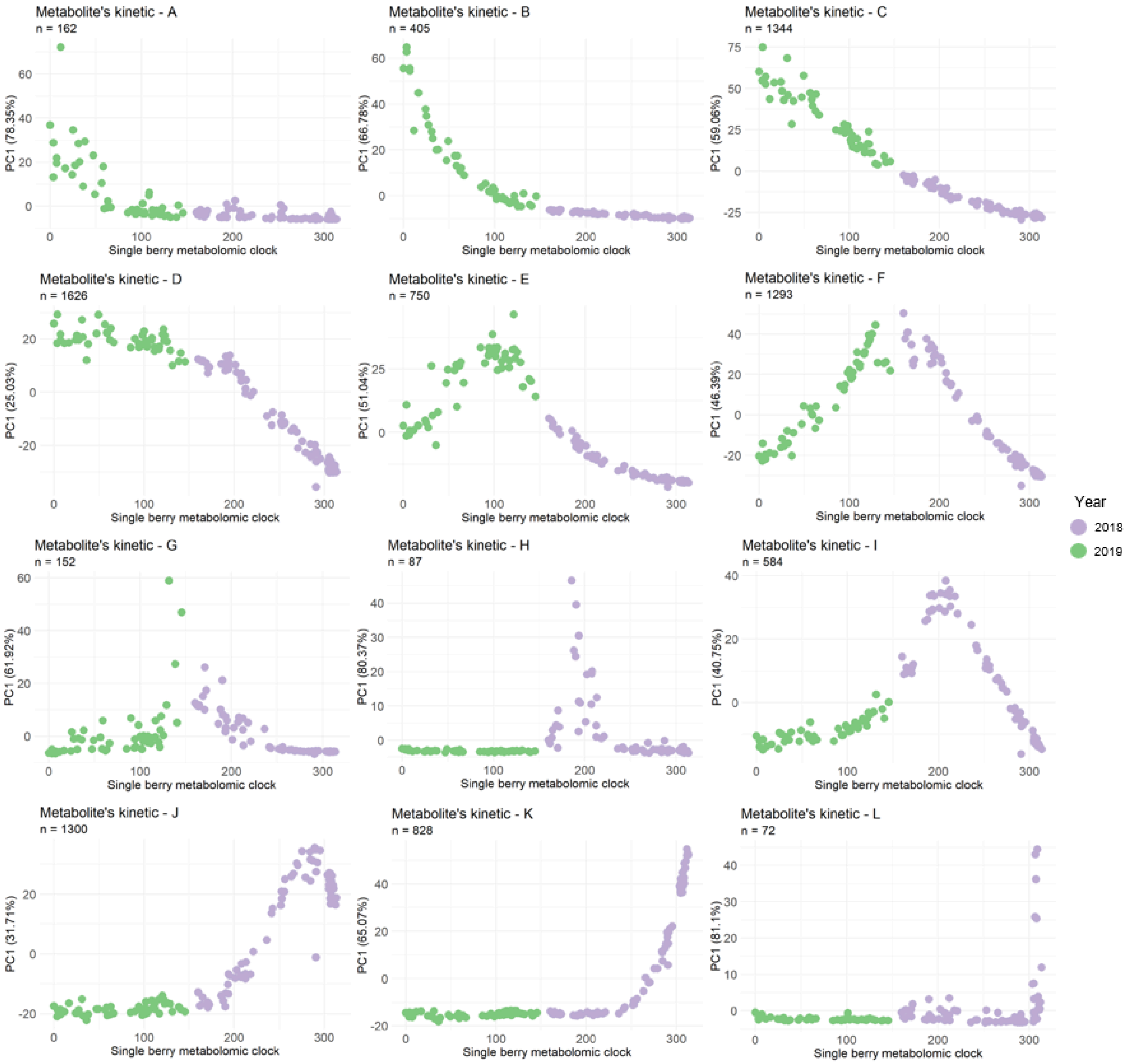


**Figure S4. Clusters representative profiles colored by the year of sampling.**

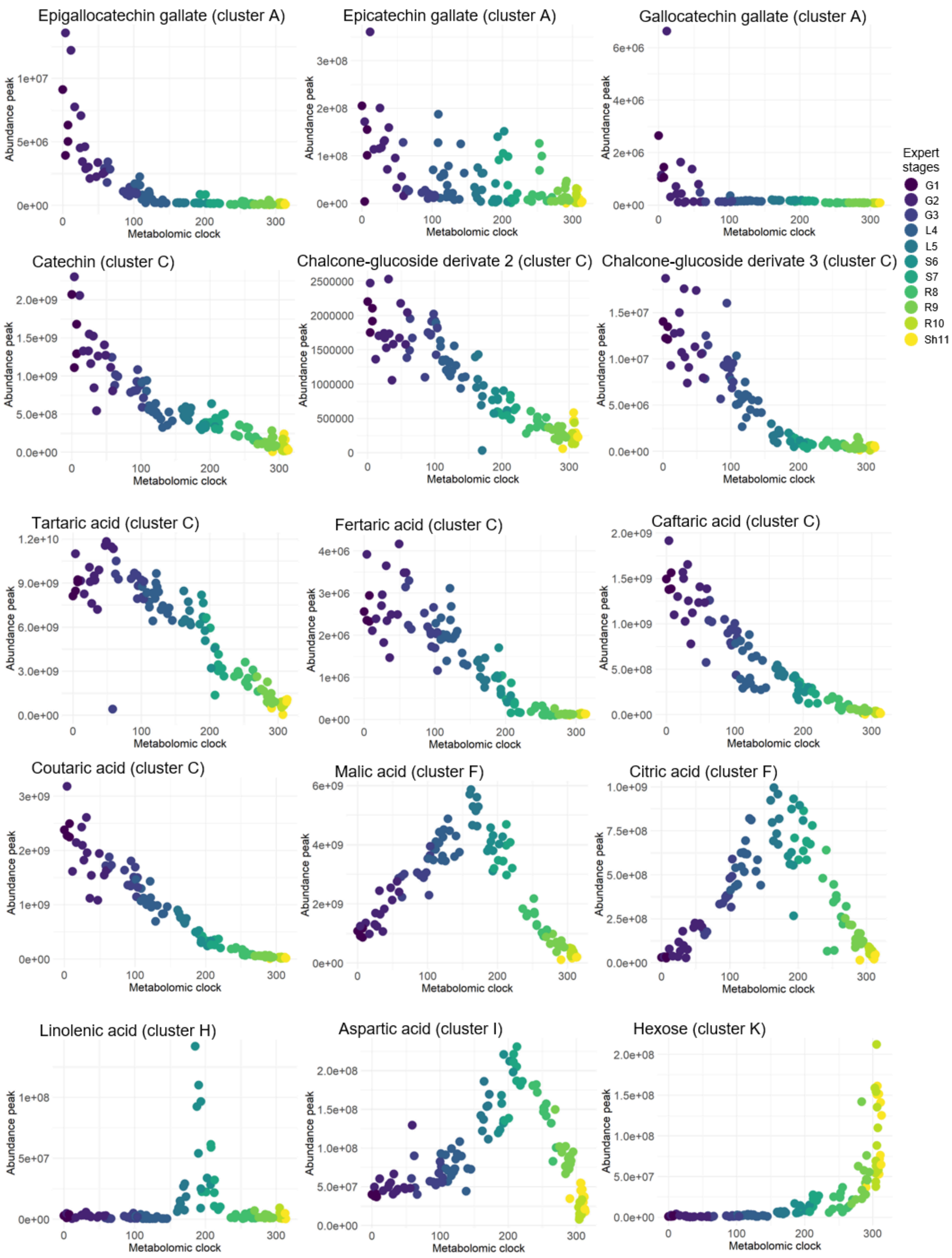


**Figure S5. Detailed profiles of the metabolites identified and presented in their pathway (**Fig. 7**), with their respective clusters, and their correlation with the cluster's eigen-metabolite.**

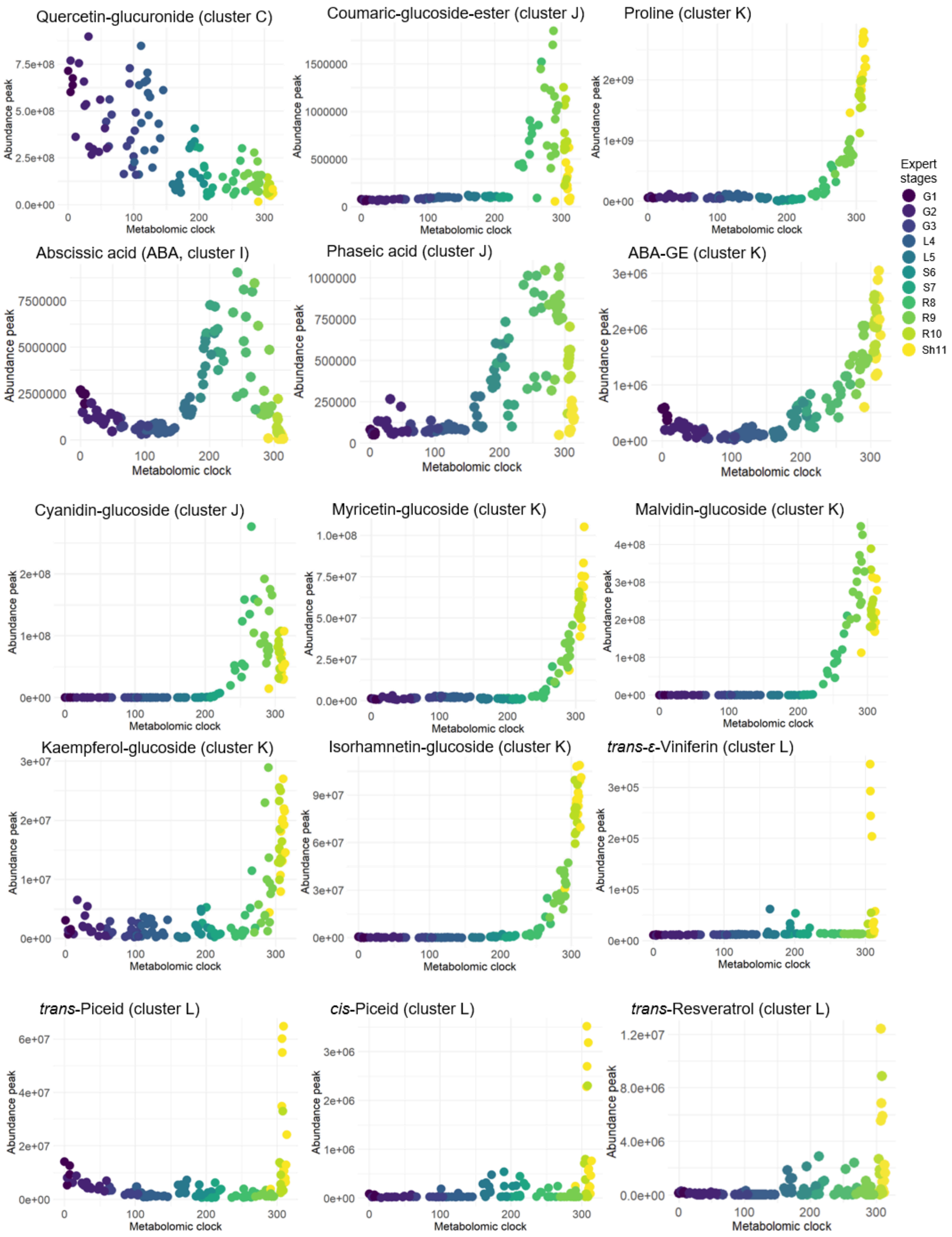


**Figure S5 *bis*. Detailed profiles of the metabolites identified and presented in their pathway (**Fig. **7), with their respective clusters, and their correlation with the cluster's eigen-metabolite.** Abscisic acid - glucoside = ABA-GE.

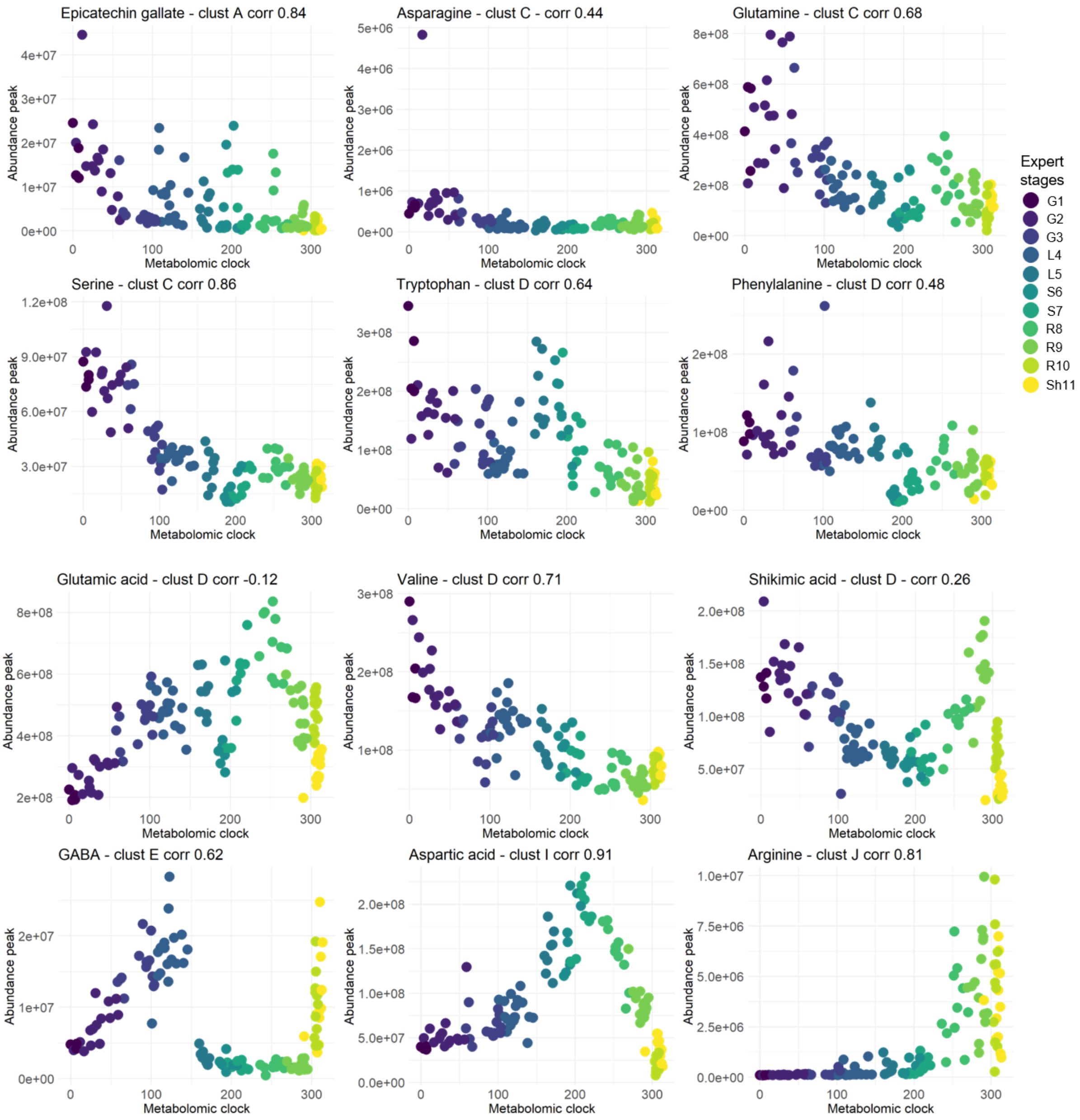


**Figure S6. Molecular standards profiles, with their respective clusters, and their correlation with the cluster's eigen-metabolite.**

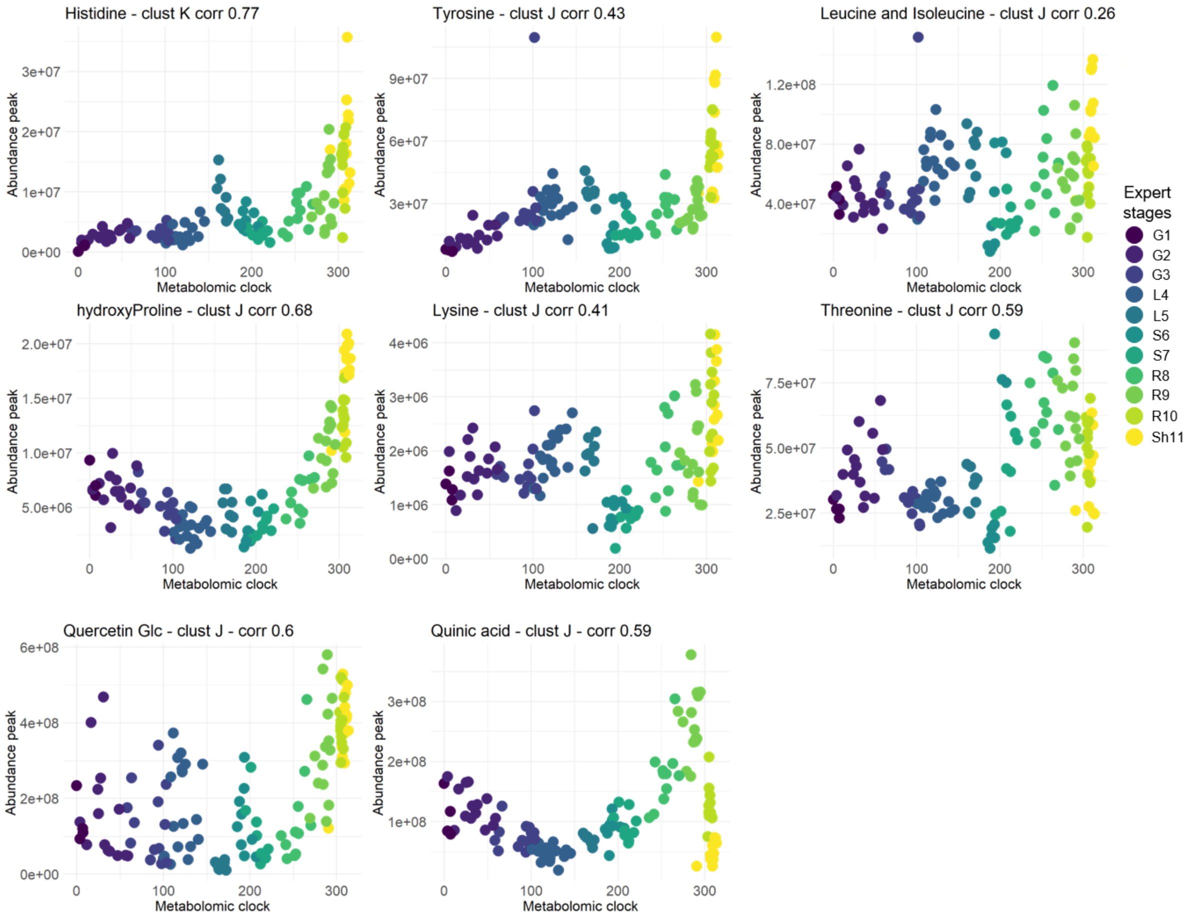


**Figure S6 *bis*. Molecular standards profiles, their respective clusters, and their correlation with the cluster's eigen-metabolite.**
